# Locus coeruleus activity improves cochlear implant performance

**DOI:** 10.1101/2021.03.31.437870

**Authors:** Erin Glennon, Angela Zhu, Youssef Z. Wadghiri, Mario A. Svirsky, Robert C. Froemke

## Abstract

Cochlear implants are neuroprosthetic devices that can provide hearing to deaf patients^1^. Despite significant benefits offered by cochlear implants, there are highly variable outcomes in how quickly hearing is restored and perceptual accuracy after months or years of use^2,3^. Cochlear implant use is believed to require neuroplasticity within the central auditory system, and differential engagement of neuroplastic mechanisms might contribute to outcome variability^4–7^. Despite extensive studies on how cochlear implants activate the auditory system^4,8–12^, our understanding of cochlear implant-related neuroplasticity remains limited. One potent factor enabling plasticity is the neuromodulator norepinephrine from the brainstem locus coeruleus. Here we examined behavioral responses and neural activity in locus coeruleus and auditory cortex of deafened rats fitted with multi-channel cochlear implants. Animals were trained on a reward-based auditory task, with considerable individual differences of learning rates and maximum performance. Photometry from locus coeruleus predicted when implanted subjects would begin responding to sounds and longer-term perceptual accuracy, which were augmented by optogenetic locus coeruleus stimulation. Auditory cortical responses to cochlear implant stimulation reflected behavioral performance, with enhanced responses to rewarded stimuli and decreased distinction between unrewarded stimuli. Adequate engagement of central neuromodulatory systems is thus a potential clinically-relevant target for optimizing neuroprosthetic device use.

---

Cochlear implants are major biomedical devices, and an exemplary success story of the application of foundational neuroscience research and use of brain-machine interface neuroprosthetics to treat a widespread neurological condition: hearing loss^1–5^. However, the auditory benefits provided by a cochlear implant are not instantaneous, in contrast to the amplification of acoustic input provided by commercial hearing aids. Some patients acquire a degree of speech comprehension with the cochlear implant a few hours after activation, but many patients unfortunately require months or even years post-implantation to achieve optimum levels of speech perception^2,3^. There are many open questions about the behavioral characteristics of this adaptation process in human listeners and the underlying neurophysiological changes^13,14^. Measuring how cochlear implants activate the central auditory system or other brain areas is technically complicated due to significant limitations with imaging in patients with implanted metallic medical devices^15^. Historically there have also been considerable challenges with experimental animal models of cochlear implant use, especially with the aims of monitoring and manipulating neural activity in implanted freely-behaving subjects. Here we addressed these issues by utilizing our recently-developed system for studying behaviorally- and physiologically-validated cochlear implant use in rats^16^, and examined neuromodulation and plasticity for cochlear implant learning and performance.

## Cochlear implant outcome variability

We initially aimed to determine how rapidly hearing could be restored to profoundly deafened rats with cochlear implants. We trained normal-hearing rats on an auditory self-initiated go/no-go task^16–18^. Animals were acoustically trained prior to deafening and cochlear implantation, in order to separate aspects of procedural task structure learning from stages of perceptual learning with the cochlear implant. In this task, rats were trained to self-initiate trials via nosepoke (**Fig. 1a**, **Extended Data Fig. 1**), and a tone of a given frequency was presented (0.5-32 kHz at one octave spacing, 100 msec duration). Rats then had opportunity to respond to a specific target frequency (4 kHz) for food reward, but were trained to withhold responses to non-target foil tones (0.5, 1, 2, 8, 16, and 32 kHz). Performance on this task was assessed by the discriminability index d’, which is the difference in z-scores for responses to targets (‘hits’) versus responses to foils (‘false positives’). Here, d’ = 0.0 indicates chance performance, while d’ ≥ 1.0 indicates good discriminability.

**Figure 1.**
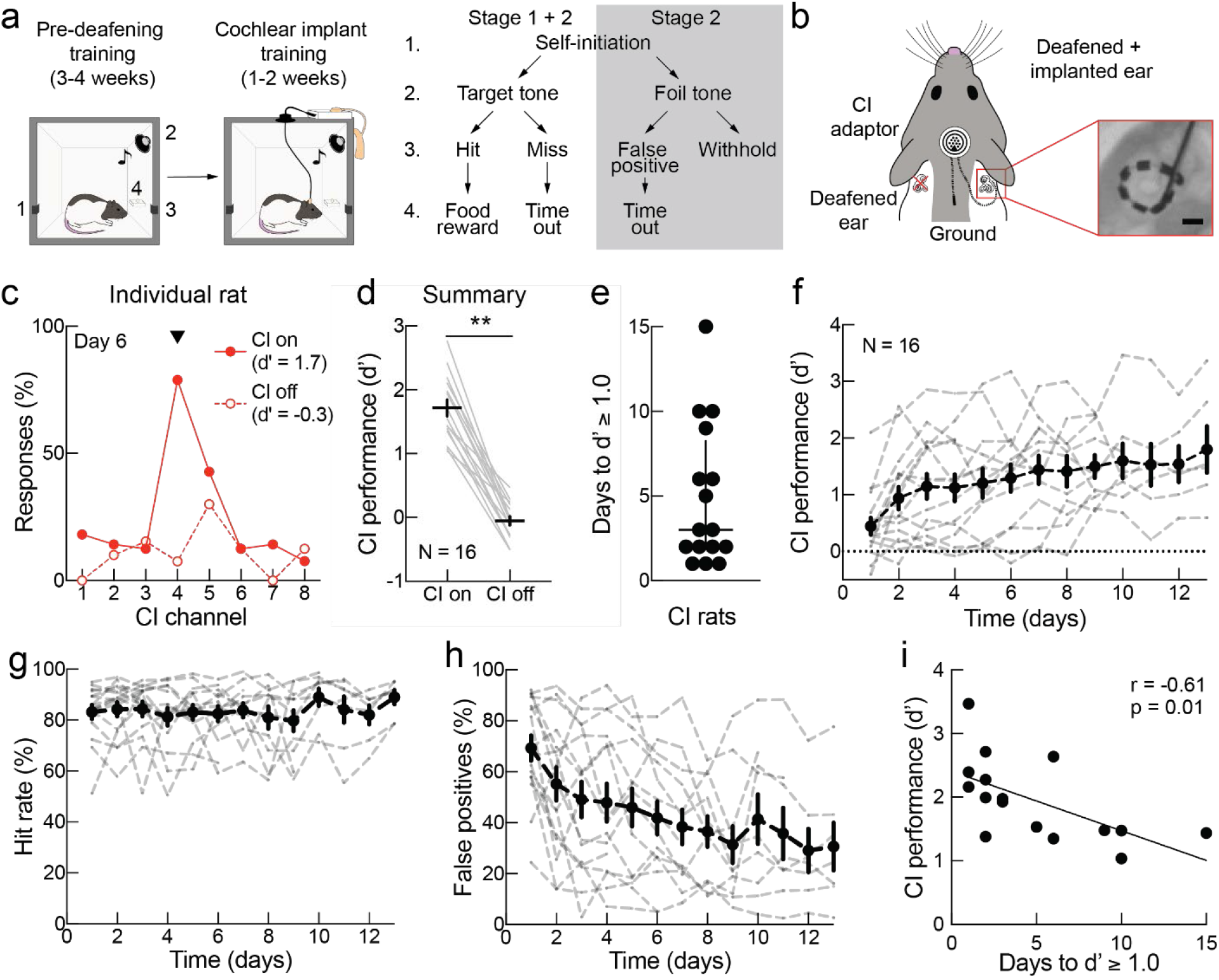
Behavioral assessment of cochlear implant learning. **a**, Schematic of behavioral training for normal-hearing and cochlear implant behavior (left). Trial structure (right): rats self-initiate trials (1) for presentation of either target or foil pure tone (2); behavioral responses (3) lead to food rewards on target hit trials, no outcome on correct withholds, or 7 s timeouts on miss trials or false positives (4). **b**, Schematic of bilateral deafening and unilateral placement of cochlear implant (CI) in rats (left). X-ray of cochlear implant electrode within the cochlea, note full >360° turn (right). Scale bar, 1 mm. **c,** Example animal showing response rates across cochlear implant channels on day 6 of behavioral training with cochlear implant with the implant on (filled) or off (open). Arrowhead, target tone programmed to activate channel 4. **d,** Behavioral performance in rats trained with the cochlear implant; performance decreased to chance when then implant was turned off (on vs off, N = 16 rats, p < 0.0001; paired two-tailed Student’s t-test). Error bars, mean ± s.e.m. **e,** Days to d’ ≥ 1.0 across rats. Error bars, median ± interquartile range. **f,** Performance (d’) with the cochlear implant over time. Gray dashed lines, individual rats. Error bars, mean ± s.e.m. **g,** Hit rates. **h,** False positives. **i,** Days to d’ ≥ 1.0 was correlated with maximum d’ for each rat (N = 16, Pearson’s r = −0.61, p = 0.01). **, p < 0.01.

After animals reached criteria (d’ ≥ 1.0 for 5+ days), they were bilaterally deafened (via cochlear lesion) and fitted with unilateral 8-channel cochlear implants (3-8 active channels per animal)^16^. These cochlear implanted rats were then re-trained to perform this task, with acoustic stimuli activating different implant channels (**Fig. 1a-b**). In order to normalize audibility across animals, cochlear implant stimulation levels were based on evoked compound action potential (ECAP) thresholds from each active cochlear implant channel (see Methods). The target and foil tones used for the cochlear implant version of the task were adjusted in order to correspond to the number of active channels (as shown in the frequency allocation tables, **Extended Data Fig. 2a-b**). Each tone predominantly stimulated one active channel as confirmed by electrodograms (**Extended Data Fig. 2c**).

Cochlear implant training occurred in two stages, paralleling the go/no-go structure of the task. In stage one, only tones activating the target channel were presented (and rewarded upon nosepoke). In stage two, other tones activating 2-7 foil channels were introduced. As with human subjects, initial performance of cochlear implanted rats attempting to respond to acoustic percepts could be variable and was often quite low^2,3,13^. Within 15 days of training, all implanted subjects reached a d’ of at least 1.0 (**Fig. 1c-f**). Implanted animals on average responded to the tone activating the target channel and withheld responses to tones activating foil channels (d’ = 1.7 ± 0.1, mean ± s.e.m., N = 16 rats, d’ quantified between days 4-14; **Fig. 1c,d**, **Supplementary Movie 1**). To verify that animals were indeed profoundly deaf and performed the task via cochlear implant stimulation (i.e., there was no residual hearing), we turned off the cochlear implant for a subset of trials and observed that behavioral performance dropped to chance (d’ = −0.1 ± 0.1, N = 16, p < 0.0001; **Fig. 1c-d, Extended Data Fig. 3,4**).

Individual rats had different learning rates throughout stage two, as measured by time to d’ ≥ 1.0 (time = 3.0 ± 6.3 days, median ± interquartile range; **Fig. 1e**). Whereas some animals seemed to quickly recognize the behavioral meaning of cochlear implant stimulation in ≤ 3 days (N = 9), other animals took 5-15 days (N = 7; **Fig. 1e-f**). This subject performance variability was not explained by differences in insertion depth, impedance, ECAP thresholds, or normal-hearing performance across animals (**Extended Data Fig. 5a-f**), similar to observations in human studies^13^. Hit rates were also comparable across animals regardless of learning rate (**Fig. 1g, Extended Data Fig. 5g**). Instead, we noticed that improvements in performance were mainly driven by a decrease in false positive rates over days (**Fig. 1h**), which was significantly slower in under-performing animals (p = 0.01, **Extended Data Fig. 5h**). The overall performance obtained by each cochlear implanted animal was inversely related to the days to reach d’ ≥ 1.0 (Pearson’s r = −0.61, p = 0.01; **Fig. 1i**), which was not the case when these same animals initially learned the task when they were normal-hearing (**Extended Data Fig. 5i**). These results indicate that after cochlear implantation, early performance predicts peak performance, similar to reports in human cochlear implant subjects^19^.

## Locus coeruleus activity during cochlear implant learning

We aimed to understand the central mechanisms contributing to individual variation in learning rates. One region important for early perceptual learning is the brainstem locus coeruleus, which is thought to broadcast a noradrenergic arousal signal throughout the central auditory system including auditory cortex^17,20–23^. To ask how locus coeruleus noradrenergic neurons might be activated during cochlear implant learning, we performed fiber photometry in tyrosine hydroxylase (TH)-Cre rats that had viral expression of GCaMP6s with AAVDJ-ef1a-DIO-GCaMP6s targeted unilaterally to locus coeruleus (**Fig. 2a-b**). We initially confirmed correct fiber placement and quality of fiber photometry recordings using tail pinch to elicit locus coeruleus responses^24^ (**Fig. 2c**). As animals were first conditioned just with target tones (in stage one) and foil tones added later (in stage two), we studied the responses in these two stages separately.

**Figure 2.**
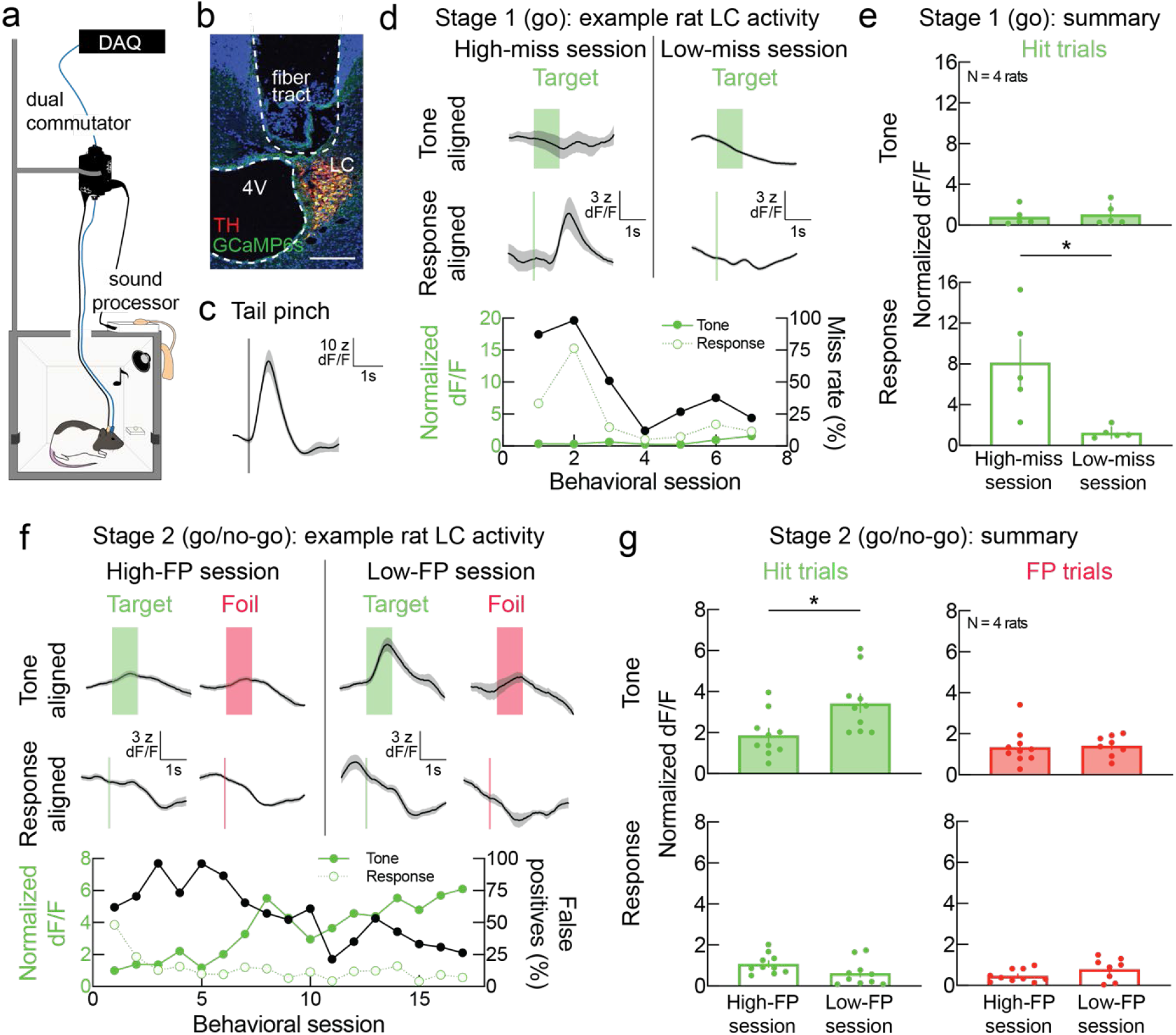
Dynamic locus coeruleus activity during cochlear implant learning. **a,** Schematic of fiber photometry recordings in rats with cochlear implants (DAQ, data acquisition system). **b,** GCaMP6s (green) in TH+ (red) cells of the locus coeruleus (LC), with fiber tract above LC (4V, 4^th^ ventricle). Scale bar, 200 µm. **c,** Locus coeruleus activity recorded in response to noxious stimulus. **d,** Example locus coeruleus activity during stage one training with cochlear implant, either aligned to tone onset (left) or behavioral response (right). Behavioral response-aligned locus coeruleus activity, tone-aligned locus coeruleus activity, and miss rate across time (bottom). **e,** Tone-aligned normalized dF/F in locus coeruleus was similar in behavioral sessions with high miss rates (bottom quartile) and behavioral sessions with low miss rates (top quartile) (top, unpaired two-tailed Student’s t-test, p = 0.71, N = 4 rats, n = 21 behavioral sessions). Response-aligned normalized dF/F in locus coeruleus was higher during behavioral sessions with high miss rates than in behavioral sessions with low miss rates (bottom, unpaired two-tailed Student’s t-test, p < 0.02). **f,** Example locus coeruleus activity during stage two (foil and target training). Hit and false positive (FP) trials in behavioral session with high false positives (left). Hit and false positive trials in behavioral session with low false positives (right). Trial-averaged locus coeruleus activity aligned to tone onset (top) or to behavioral response (middle). Behavioral response-aligned locus coeruleus activity, tone-aligned locus coeruleus activity, and false positives across time (bottom). **g,** Tone-aligned normalized dF/F in locus coeruleus on hit trials was higher in behavioral sessions with low false positives (top quartile) than in behavioral sessions with high false positives (bottom quartile) (top left, unpaired two-tailed Student’s t-test, p = 0.01, N = 4 rats, n = 40 behavioral sessions), but not on false positive trials (top right, unpaired two-tailed Student’s t-test, p = 0.85). Response-aligned normalized dF/F in locus coeruleus was similar between behavioral sessions with high and low false positives both hit trials (bottom left, unpaired two-tailed Student’s t-test, p = 0.08) and false positive trials (bottom right, unpaired two-tailed Student’s t-test, p = 0.12). Data are mean ± s.e.m. *, p < 0.05.

We quantified the trial-averaged locus coeruleus responses to the stimulus (‘tone-aligned’), and responses aligned to the nosepokes (‘response-aligned’) which immediately precede reward dispensation on hit trials. We found that locus coeruleus activity was not fixed, but instead dramatically changed over the course of cochlear implant learning. Specifically, locus coeruleus activity shifted from being driven by unexpected reward (in stage one) to being evoked by target stimulus presentation (in stage two), predicting improvements across cochlear implant subjects for when each animal reduced miss rates in stage one and began to have lower false positives in stage two.

The first effect we observed is represented by the example responses shown in **Figure 2d**. Task-related locus coeruleus responses emerged just prior to when this animal first began using the cochlear implant to respond to the target tone during stage one. These initial locus coeruleus responses were linked to reward and behavioral response, without clear stimulus-evoked signals (**Fig. 2d, left, bottom**). During this first stage of training, once this rat began responding consistently to the target tone with lower miss rates, locus coeruleus activity suddenly decreased (**Fig. 2d, right, bottom**). Across animals and sessions, there was negligible target tone-aligned locus coeruleus activity during earlier high-miss sessions and later low-miss sessions, both for the hit trials (**Fig. 2e, top**; N = 4 rats, n = 21 behavioral sessions split into quartiles, p = 0.71 comparing top versus bottom quartiles, unpaired two-tailed Student’s t-test) and for miss trials (**Extended Data Fig. 6a,b**). In contrast, there was substantial response-aligned locus coeruleus activity during earlier high-miss sessions, which decreased as animal performance increased (**Fig. 2e, bottom**; p < 0.02).

Animals were then moved to stage two of training when foil tones were also presented. In the animal shown in **Figure 2f** (same rat as **Fig. 2d**) during stage two training, when false positive rates were high during earlier sessions, the tone-aligned and response-aligned locus coeruleus activity both remained low (**Fig. 2f, left, bottom**). As this animal began to learn to withhold responses to foil tones leading to lower false positive rates, we observed target tone-aligned locus coeruleus activity that preceded reward (**Fig. 2f, right, bottom**). Across animals, tone-aligned locus coeruleus activity was higher on hit trials throughout behavioral sessions with low false positive rates compared to behavioral sessions with high false positive rates (**Fig. 2g left, top**; N = 4 rats, n = 40 behavioral sessions divided into quartiles, p = 0.01 comparing top versus bottom quartiles). There was negligible response-aligned activity (**Fig. 2g, left, bottom**; p = 0.85), and during false positive, miss, and withhold trials, locus coeruleus activity was similarly low (**Fig. 2g, right, Extended Data Fig. 6c-d**).

Plasticity of locus coeruleus activity seemed to reflect changes to internal representations of task variables and was not modality specific just for cochlear implant use. We also performed fiber photometry from locus coeruleus in normal-hearing TH-Cre animals trained on the acoustic version of this go/no-go task (**Extended Data Fig. 7**). We observed similar patterns and dynamics of locus coeruleus activity when the target tone was changed from 4 kHz to a different frequency. These results indicate that the locus coeruleus neuromodulatory signal is initially driven by unexpected reward, but becomes linked to stimuli predicting this reward over the course of training.

## Locus coeruleus stimulation accelerates cochlear implant learning

We next wondered if we could harness the activity of the locus coeruleus to enhance learning with the cochlear implant. The imaging studies of **Figure 2** demonstrate that once animals learned the task, target tones selectively activated locus coeruleus on correct trials. We therefore hypothesized that pairing locus coeruleus activation with presentation of the target tone earlier in training could accelerate cochlear implant learning, analogous to the effects of locus coeruleus stimulation on enhanced auditory cortical representations and perceptual learning in normal-hearing rats^17^.

Locus coeruleus was stereotaxically targeted in each animal and identified via electrophysiological responses in vivo to toe pinch (**Extended Data Fig. 8a,b**). We then expressed the excitatory opsin ChETA in noradrenergic locus coeruleus neurons via injection of AAV5-ef1a-DIO-ChETA in acoustically trained TH-Cre rats. We confirmed ChETA expression in TH+ locus coeruleus cells using immunohistochemistry and examined placement of the optical fiber within locus coeruleus via micro-magnetic resonance imaging/micro-computed tomography (µ-MRI/µ-CT) co-registration performed by blinded observers (**Fig. 3a, Extended Data Fig. 8c**). Viable animals with mistargeted optical fibers were used as sham controls. Animals were deafened and fitted with a cochlear implant two weeks after injection, and behavioral training began several days later. Starting on the first day of cochlear implant training, optogenetic stimulation of locus coeruleus was paired with the target tone for 5-10 minutes prior to each session (‘offline LC pairing’, **Fig. 3b**). We conducted pairing outside of behavioral context to focus on the impact of potential longer-term central modifications to neural circuits induced by locus coeruleus pairing, rather than more immediate changes to arousal level or brain state triggered by noradrenergic modulation.

**Figure 3.**
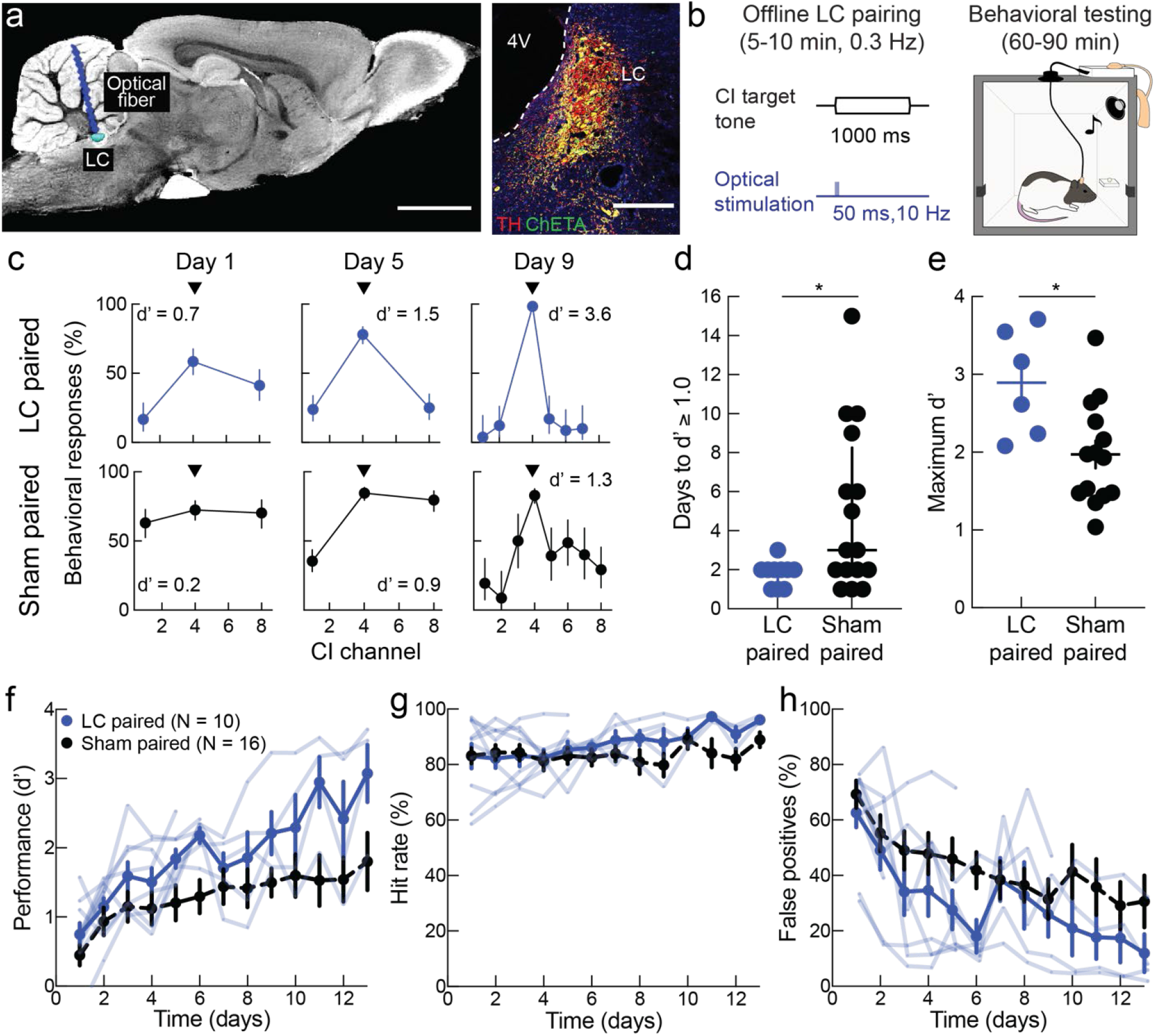
Locus coeruleus pairing enhances cochlear implant learning. **a,** Optogenetic stimulation of noradrenergic locus coeruleus. Left, optical fiber placement in locus coeruleus (LC) was confirmed by µ-CT/µ-MRI co-registration. Scale bar, 3 mm. Right, ChETA expression (green) in TH+ cells (red) in locus coeruleus. Scale bar, 200 µm. **b,** Schematic of offline locus coeruleus pairing and cochlear implant (CI) training. **c,** Example animals receiving either locus coeruleus pairing (top) or sham pairing (bottom), showing behavioral responses and d’ on days 1, 5, and 9 of cochlear implant training. Arrowhead, target tone programmed to activate channel 4 in both animals. Error bars, 95% confidence intervals. **d,** Locus coeruleus paired rats reach criteria (d’ ≥ 1.0) more quickly than sham paired rats (locus coeruleus paired, N = 10 rats vs sham paired, N = 16 rats, p = 0.04, two-tailed unpaired Mann–Whitney test). Error bars, median ± interquartile range. Note that all paired animals reached criterion within 1-3 days. **e,** Locus coeruleus paired rats with at least six days of cochlear implant training had higher maximum d’ than sham paired rats (locus coeruleus paired, N = 6 rats vs sham paired, N = 14 rats, p = 0.01, two-tailed unpaired Student’s t-test). Error bars, mean ± s.e.m. **f,** Implant performance (d’) over time in locus coeruleus paired rats vs in sham paired rats. **g,** Hit rates. **h,** False positives. Error bars, mean ± s.e.m. Four locus coeruleus paired rats and two sham paired rats did not reach the six day performance requirement to calculate maximum d’, these animals are displayed in **d**,**f-h** but excluded from **e**. *, p < 0.05; **, p < 0.01.

Remarkably, all animals receiving locus coeruleus pairing reached d’ ≥ 1.0 within three days, i.e., they learned to use the cochlear implant more quickly compared to ‘sham paired’ animals injected with control YFP virus (AAV5-ef1a-DIO-eYFP) or off-target fiber implantation (**Fig. 3c,d, Extended Data Fig. 9a**; locus coeruleus paired animals: 2.0 ± 1.0 days to d’ ≥ 1.0, median ± interquartile range; sham paired animals: 3.0 ± 6.3 days to d’ ≥ 1.0; Mann-Whitney test, p = 0.04). Locus coeruleus paired animals also had higher levels of maximum performance (**Fig. 3e,f, Extended Data Fig. 9b-e**; locus coeruleus paired animals: d’ = 2.9 ± 0.3, mean ± s.e.m; sham paired animals: d’ = 2.0 ± 0.2; unpaired two-tailed Student’s t-test, p = 0.01). Enhanced learning in locus coeruleus paired animals was unrelated to differences in cochlear implant functionality assessed by insertion depth, impedance, ECAP levels, or behavioral engagement in the task (**Extended Data Fig. 10a-e**). Performance in locus coeruleus paired animals improved over time (**Fig. 3f**), with behavioral gains driven by a sustained high hit rate and progressive reduction in false positive rates similar to the changes in behavior in sham paired animals (**Fig. 3g,h**).

We conclude that outcome variability across individual cochlear implant users was not entirely due to differences in insertion quality or level of peripheral auditory system activation. Specifically, none of these factors were different across the groups of cochlear implant rats examined here (**Extended Data Figs. 5,10**). Instead, variable learning rates across individuals seem to be due to differential engagement of modulatory systems (e.g., locus coeruleus) known to promote long-lasting plasticity and enhance perceptual learning. Although we artificially augmented neuromodulatory activity with optogenetic stimulation in locus coeruleus paired animals, our results from **Figure 2** and **Extended Data Figure 7** show that this system is endogenously active during acoustic or cochlear implant training, perhaps at different levels in various individuals. Such a continuum of neuromodulatory engagement might then account for behavioral variability across the population of cochlear implant users.

## Cortical representations of cochlear implant stimuli

The locus coeruleus projects throughout the central auditory system including the auditory cortex, suggesting that differing degrees of noradrenergic modulation might lead to variable neural representations of cochlear implant channels. We focused on neural responses to implant stimulation in the auditory cortex, based on evidence suggesting that behavioral improvements with cochlear implants are paralleled by changes in auditory cortical responsiveness^10,25–27^. We specifically asked how cortical responses might relate to the behavior of rats trained with cochlear implants either with or without locus coeruleus stimulation. Locus coeruleus pairing with auditory stimuli in normal-hearing rats leads to lasting changes in auditory cortex but not the auditory thalamus^17,20^, and auditory cortical activity is required to perform this acoustic go/no-go task^28^.

We performed electrophysiological recordings in anesthetized rats after these animals had been deafened and trained to use cochlear implants; some animals were locus coeruleus paired while other animals were sham paired. Auditory cranial nerve (‘CN VIII’) ECAPs were measured and multi-unit recordings from auditory cortical responses to each individual cochlear implant channel were recorded (**Fig. 4a**). These methods are comparable to the types of electrophysiological recordings that can be conducted in human subjects; ECAPs are routine in human users, and multi-unit activity reflects a population-level responsiveness similar to electrocorticography and electroencephalography but with higher spatial resolution and signals closer to underlying single-unit representations.

**Figure 4.**
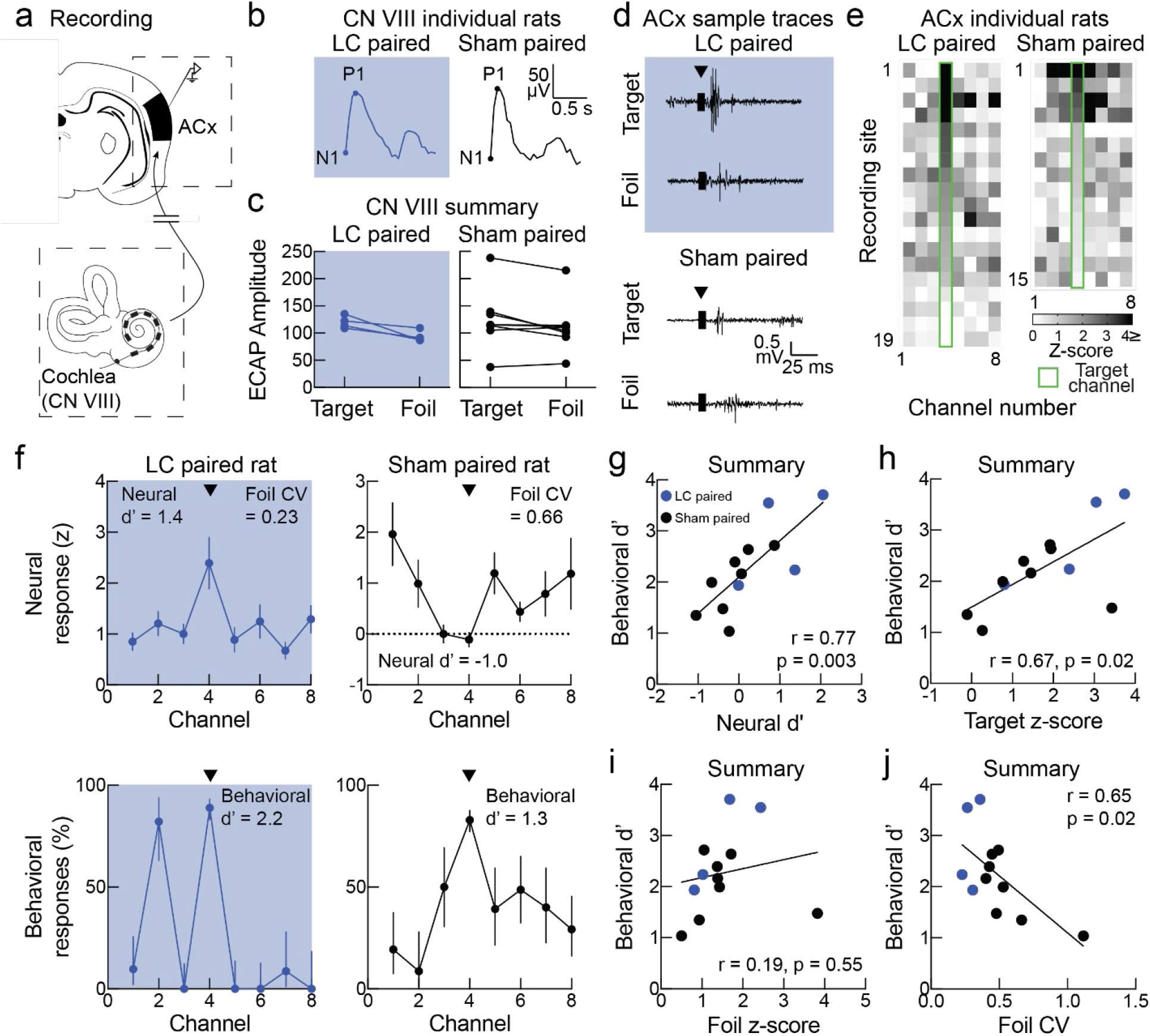
Auditory cortical responses in cochlear implant trained rats. **a,** Schematic of recording sites in the auditory pathway. **b,** Example ECAPs in CN VIII from a locus coeruleus (LC) paired rat (left) and a sham paired rat (right). **c,** Average ECAP amplitude (P1-N1) did not differ across group and channel valence (target vs foil, sham vs locus coeruleus, p = 0.70; one-way ANOVA with Bonferroni correction). **d,** Example traces of CI evoked auditory cortical responses from target or foil channels in a locus coeruleus paired and a sham paired rat. Arrowhead, cochlear implant stimulus artifact (amplitude clipped for display purposes). **e,** CI evoked auditory cortical responses across CI channels and recording sites in a locus coeruleus paired rat (left) and a sham paired rat (right). Green outline indicates behavioral CI target channel. **f,** Neural response curves (top) and behavioral responses curves (bottom) for a locus coeruleus paired animal (left) and a sham paired animal (right). Error bars, mean ± s.e.m. (top) or 95% confidence intervals (bottom). **g,** Across locus coeruleus paired (N = 4, blue circles) and sham paired animals (N = 8, black circles), neural d’ correlated positively with behavioral d’ (N = 12, Pearson’s r = 0.77, p = 0.0003). **h,** Normalized neural responses to the target channel correlated positively with behavioral performance (Pearson’s r = 0.67, p = 0.02). **i,** Normalized average responses to the foil channels did not correlate with behavioral performance (Pearson’s r = 0.19, p = 0.55). **j,** The coefficient of variance of neural responses to foil channels negatively correlated with behavioral performance (Pearson’s r = 0.65, p = 0.02).

Auditory nerve ECAPs were comparable across target and foil channels as well as across groups (**Fig. 4b-c**). This indicates that overall auditory nerve activity was broadly similar for both groups and stimulus types. In contrast, we noticed that the cortical multi-unit responses to target channel stimulation seemed higher on average in locus coeruleus paired animals versus sham paired animals (**Fig. 4d-f**). Based on this observation, we calculated neural d’ values (mean target z-score – mean foil z-score across all multi-unit recording sites) for each animal and compared this to their behavioral performance. We found that neural and behavioral d’ values were highly correlated across both groups of animals, such that animals (either sham or locus coeruleus paired) with high behavioral performance also had higher neural d’ values (Pearson’s r = 0.77, p = 0.003; **Fig. 4g**).

This correlation was due to two main indicators of enduring cortical plasticity resulting from cochlear implant training (augmented in some animals by explicit locus coeruleus pairing). These effects can be observed in the example animal with locus coeruleus pairing shown in **Figure 4f** (left), which had good performance using the cochlear implant to perform the task (d’ = 2.2); recordings from this animal showed 1) strong cortical responses to the target channel, and 2) very similar responses across foil channels. In contrast, a sham paired animal with lower performance had more variable responses over all channels (**Figure 4f**, right). Across animals, behavioral performance was correlated with responses to the target channel but not foil channels (targets: Pearson’s r = 0.67, p = 0.02; **Fig. 4h**; foils: Pearson’s r = 0.19, p = 0.55; **Fig. 4i**). Instead of the mean response to foils, behavioral outcomes were inversely correlated with the variability of foil channel responses among the different active channels (as quantified by the coefficient of variation, the average standard deviation across foils normalized by the average mean response; Pearson’s r = 0.65, p = 0.02; **Fig. 4j**). The reduction in variation means that the different foil channels have similar evoked responses to each other, over the active channels and throughout the multi-unit recording sites from each animal. Therefore, cochlear implant stimulation can effectively engage the auditory cortex in trained rats, with cortical responses shaped via mechanisms of neuromodulator-enabled plasticity to represent behavioral categories of different input. Specifically, reward-predictive stimuli evoked stronger neural responses, whereas responses to unrewarded stimuli seemed to be grouped together and became less distinct from each other.

For enhancement of perceptual and cognitive abilities, adequate interface between neuroprosthetics and neural tissue requires adaptation by the host biological circuitry to the signals provided by the neural implant. Here we discovered that locus coeruleus activity is a key indicator of hearing restoration with clinical-grade multi-channel cochlear implants in profoundly deaf rats. Responses in the locus coeruleus could predict when each animal first began hearing with the implant as well as overall performance levels. Optogenetic stimulation augmented the natural engagement of locus coeruleus to normalize performance across all animals. Failure to optimally use the cochlear implant in some subjects was therefore not due to poor activation of the auditory system, and indeed we observed robust cortical responses to implant stimulation in every animal. Training and neuromodulation seemed to refine cortical activity in a manner related to the behavioral meaning of the sounds, leading to an effective perceptual categorization in auditory cortex of the best performing animals. Our results provide a potential path for accelerating and improving outcomes with cochlear implants as well as other neuroprosthetic devices. Targeted neuroplasticity combined with neuromodulatory recruitment could be achieved with devices such as vagus nerve stimulators (believed to activate locus coeruleus)^29,30^, combined with real-time behavioral monitoring to enhance prosthetic device use during optimal periods of attention and arousal.

## Methods

### Ethics

All procedures were approved under NYU Langone Institutional Animal Care and Use committee protocols.

#### Surgical protocols

All animals were kept in a vivarium on a 12/12-hour light/dark cycle and housed individually or in pairs. Female Long-Evans wild-type or TH-Cre rats 3-5 months old were anesthetized with intramuscular ketamine (40 mg/kg) and dexmedetomidine (0.125 mg/kg). Atropine (0.02 mg/kg) and dexamethasone (0.2 mg/kg) were subcutaneously applied immediately after anesthesia induction to minimize bronchial secretions, inflammation, and intracranial pressure. Body temperature was maintained slightly hypothermic at 34-35°C with a DC Temperature Controller heating pad throughout the procedure and eye ointment (Puralube® Vet Ointment, Dechra) used to prevent corneal drying. The rat was positioned prone to optimize respiratory function.

#### Viral injections and optic fiber placement in locus coeruleus

Viral injections into female rat locus coeruleus^18^ were performed using stereotaxic coordinates (from lambda, in mm: 3.7-3.8 posterior, 1.2-1.4 lateral, 5.6-6 ventral) with the head at a 15° downward angle. A craniotomy was placed over the left locus coeruleus and location was verified during surgery by measuring multi-unit spontaneous activity and responses elicited by noxious stimuli (tail pinch) and confirmed afterwards using histological methods and µ-CT/µ-MRI co-registration (**Extended Data Fig. 6**). Injections were performed with a 5 µL Hamilton syringe and a 33-gauge needle. We used two 1.0 µl injections at 0.1 nl/s into locus coeruleus: one at the most dorsal point where noxious stimuli responses were observed and the second 250 µm ventral to that location.

For viral expression in noradrenergic locus coeruleus neurons, we used TH-Cre Long-Evans rats to restrict expression of Cre-inducible viruses^31^. For optogenetic stimulation, we injected AAV5-ef1α-DIO-ChETA-EYFP (viral titer: 1.5×10^13^); for sham stimulation we injected AAV5-ef1α-DIO-EYFP (viral titer: 1.0×10^13^); and for fiber photometry we injected AAVDJ-ef1α- DIO-GCaMP6s (viral titer: 1.1×10^13^, Stanford Neuroscience Gene Vector and Virus Core, Deisseroth lab).

A calibrated optical fiber ferrule (200 µm for optogenetic experiments, 400 µm for fiber photometry experiments, ThorLabs) was implanted in locus coeruleus, and craniotomy and implant were sealed with silicone sealant and Metabond® (Parkell Inc).

#### Deafening and cochlear implantation

Cochlear implantation was conducted as previously described^16^; for TH-Cre rats, this occurred 2-4 weeks after viral injection. For both deafening and cochlear implantation, a postauricular incision was made and the superficial fascia of the neck was dissected. The sternocleidomastoid muscle and posterior belly of the digastric muscle were retracted from the tympanic bulla. The tympanotomy was begun ventrocaudally to the trunk of the facial nerve with a 0.5-mm diamond burr and continued dorsally until the stapedial artery overlying the round window was fully visualized.

The cochleostomy site was identified ∼0.5 mm directly below the lip of the round window in the basal turn of the cochlea. The site was drilled with a 0.1-mm diamond burr. For the deafened only side, a sham cochlear implant array (4 channel, Cochlear Ltd.) was inserted and left in the cochlea for 10 minutes to induce full field deafness. The array was removed and the cochleostomy site is closed with a tympanic bulla muscle graft followed by 2-octyl cyanoacrylate (Surgi-Lock 2oc, Meridian™ Animal Health). For the cochlear implanted side, prior to performing the cochleostomy and inserting the array (8 channel, Cochlear Ltd.), the array lead and connector were cemented to the head cap securing the previously implanted optic fiber. Blunt dissection was used to expand the postauricular incision towards the head cap. The ground was sutured into a small muscle pocket in the trapezius. The array was inserted without resistance using AOS forceps (Cochlear Ltd.) until all the platinum-iridium contacts were within the scala tympani. The array and cochleostomy were sealed using 2-octyl cyanoacrylate. Before closure, dexamethasone was applied to the root of the facial nerve to prevent inflammation. The portion of the lead wires exiting the skull connector as reinforced with additional silicone insulation, and further protected with silicone sealant (Kwik-Cast, WPI) and dental epoxy (Triad Gel, Dentsply).

#### Acoustic training

Rats were trained on a go/no-go task^16–18,28,32^ to nosepoke in response to a target tone frequency for a sugar pellet reward (Bio-Serv) in operant conditioning chambers (Med Associates, Inc.). Each chamber contained a speaker calibrated across frequencies at 70 dB SPL, a food dispenser on the left wall and three nosepoke ports (two on either side of the food dispenser and one on the wall opposite). Each chamber was placed in a larger wood enclosure and insulated with foam. The measured background noise in each chamber was <30-40 dB SPL.

A total of thirty-two adult female Long-Evans rats were used for these studies. Animals were food restricted to maintain the weights at 80-85% of their initial pre-training weights. First, animals were shaped with two days of training to nosepoke for one food pellet. Next, rats were trained to nosepoke within 2.5 seconds after a target tone was played (stage 1: go training). When the rats had hit rates of >80%, stage 2 (go/no-go) training began, and three foil tones were introduced (2-16 kHz at one octave intervals excepting the target frequency). Animals were trained to hit rates >90%, along with false positive rates <40%. Finally, the foil tones were expanded to six total (0.5-32 kHz at one octave intervals excepting the target frequency), and animals were trained to the same criteria. Target and foil pure tones were 100 ms in duration presented in a pseudo-random order at 70 dB SPL. For correct trials, each trial ended at either the time of food pellet delivery (hit trials for targets) or 2.5 s after the tone (correct withhold trials for foils). On error trials, failure to respond (miss trials for targets) as well as incorrect responses (false positive trials for foils) were punished with a time-out of 7 s before the next trial began. Random nose pokes were punished with time-out as well. Rats self-initiated the trials by nosepoking in a different port than the ‘response’ port. After 0.5-1.5 s, either a target or foil tone was played. Behavioral performance was measured using the discriminability index d’ (the difference in the z-scores for the distribution of responses to targets and for the distribution of responses to foils). d’ values were computed as the difference in z-scores between hits and false positives: d’ = z(hit rate) – z(false positive rate).

Animals that achieved criterion behavioral performance on the baseline task with the target tone of 4 kHz underwent surgery as described above, had optical fibers chronically implanted in left locus coeruleus, and were allowed to recover for two weeks during viral expression. Animals then underwent bilateral deafening and unilateral cochlear implantation^16^. After three days of recovery, animals began behavioral training with the implant.

#### Cochlear implant programming

Impedance and evoked compound action potential (ECAP) threshold measurements are obtained using Custom Sound® EP and used for the initial programming of the sound processor as previously described^16^. In Custom Sound® Suite 4.0, ECAP thresholds were measured and used to set the dynamic range. The ECAP threshold was used as the comfort level and the threshold level was set to ∼6 dB (∼300 μA) below. The implant impedances and ECAP thresholds were checked every 3-5 days and changes to comfort and threshold levels made accordingly. The frequency allocation table (188 Hz to 7938 Hz) was distributed equally across the active channels (**Extended Data Fig. 2**). The pulse width was set to 25 μs, stimulation rate to 900 Hz, and stimulus maxima to 1 (such that only one electrode was stimulated at a time for any given acoustic input).

#### Cochlear implant training

Rat implants were connected to the sound processor via a custom commutator (Exmore) at the top of the behavior box. This connected to the implant emulator and sound processor (Freedom^TM^, Cochlear Ltd.) via a DB-25. The microphone of the sound processor was oriented toward the speaker. The speaker then played pure tones (1000 ms duration) corresponding to the center frequency for an electrode (**Extended Data Fig. 2b**). Speakers were calibrated based on electrodograms, which represent the assignment of stimulation pulse magnitudes to electrodes as a function of time. In Nucleus devices, this pulse magnitude sequence can be captured from the radio-frequency (RF) output of the sound processor (Cochlear Ltd.). The same model of sound processor programmed with the same processor settings used for behavioral testing was used for obtaining the electrodograms. Med Associates software was used to play the center frequency for each electrode across sound intensities (50-90 dB SPL, 2 dB steps) while an automated Matlab routine was used to record the RF output from the sound processor. RF output was captured using the Clinical Programming System (CPS) connected to a PC by an IF5 card (Cochlear Ltd.) and controlled with the NICCaptureClient commands built into the Nucleus Implant Communicator software package (Cochlear Ltd.). These commands record the pulse sequence represented in the RF output. The sound processor and CPS unit were placed with the microphone oriented towards the speaker in the operant conditioning box, as done during behavioral testing. The electrodograms were then analyzed, and the center frequency intensity for each electrode that generated 80% maximum pulse amplitude while minimally stimulating other electrodes was selected for behavioral training (**Extended Data Fig. 2c**).

Animals were re-trained on the behavioral task, beginning with the nosepoke training to confirm recovery and task engagement. Animals had to use the cochlear implant to respond to the center frequency of the target channel (stage 1). Once a rat reached a hit rate of >80% to the target, the two most distal channels were introduced as foils (stage 2). After an animal reached criteria of d’=1.0 on the two-foil version of the task, the center frequencies of the remaining active channels were introduced as foils. Since the activation of the cochlear implant was acoustic, it was imperative that the animal was profoundly acoustically deaf, as confirmed by performance dropping to chance with the cochlear implant turned off (**Fig. 1, Extended Data Figs. 3, 4, and 10**), and compared to the d’ in the session immediately preceding or following with the cochlear implant on. For the analysis of **Figure 1**, maximum d’ was the highest d’ value across all days of stage two after all active foil channels were used for training. For comparisons in **Figures 3, 4,** and **Extended Data Figure 9**, maximum d’ was computed only in animals that had at least six days of behavioral training to ensure comparisons with steady-state performance rather than initial learning rates.

#### Optogenetic pairing

Starting on the first day of target tone training with the cochlear implant, the target tone (center frequency of target channel) was paired with activation of the locus coeruleus with blue light. For optogenetic stimulation, locus coeruleus-tone pairing was conducted at a rate of 0.3 Hz, for 5-10 minutes daily immediately prior to behavioral testing. Optogenetic stimulation of locus coeruleus began at tone onset, tone duration was 1000 ms (same as during behavioral training), and locus coeruleus optogenetic stimulation was 500 ms duration at 10 Hz with 5 ms pulses. This pairing procedure was performed each day throughout training with the cochlear implant. Pairing was conducted outside of the context of behavior to reduce the potential impact of locus coeruleus activation on behavioral performance, such as freezing and changes in attention^33^.

#### Fiber photometry

To perform fiber photometry, 210 Hz sinusoidal blue light (465 nm) and 330 Hz ultraviolet light (405 nm; to control for motion and photobleaching) were delivered via the optical fiber from an LED (100-200 μW) for GCaMP6s excitation (Doric). We collected the emitted light via the same optical fiber, using an integrated fluorescence mini cube (Doric) to direct emitted light to a femtowatt silicon photoreceiver (Newport), and data were recorded using a real-time processor (RZ6, TDT). The analog readout was then low-pass filtered at 10 Hz. Behavioral boxes (Med Associates) were synchronized with the fiber photometry system using TTL pulses to align nosepokes and tone onset to locus coeruleus activity. A custom dual commutator system was built to allow for simultaneous fiber photometry recording and cochlear implant behavior (Doric, Exmore).

Recordings were analyzed using custom Matlab script. Data from both channels were smoothed using a moving average. The GCaMP signal was then corrected by normalizing each channel using a least-squares regression. The dF/F was generated by subtracting the fitted 405 nm signal from the 465 nm signal in order to reduce movement and photobleaching artifacts. To compare across animals and recording sessions, signal for each trial was z-scored using the 3 seconds prior to trial initiation. Behavioral sessions with more than 50 trials were binned into separate 50 trial blocks for analysis. Traces were trial-averaged based on behavioral response: hits, false positives, misses, or withholds. All behavioral sessions were divided into quartiles based on miss rate (stage one) or false positive rate (stage two). The neural activity from the top and bottom quartiles was compared.

#### Micro-Computed Tomography Imaging: Locus coeruleus and cochlear implant targeting confirmation

At the end of behavioral experiments, animals were scanned on CT to confirm that the cochlear implant was intracochlear and for co-registration analysis of optical fiber placement for optogentic animals. The localization of the implanted optical fibers was assessed in vivo using μ-CT scans in post-implanted rats, followed by co-registration with a µ-MRI rat brain atlas (**Extended Data Fig. 8**). Unlike µ-MRI, μ-CT imaging can be performed on subjects with metal implants. We combined registration of post-implant μ-CT with a three-dimensional µ-MRI rat brain atlas (as lack of soft tissue contrast in μ-CT limits the anatomical detail required to precisely verify optical fiber placement). The three-dimensional µ-MRI rat brain atlas was established by acquiring four age, sex, and strain matched ex vivo rats using T_2_*-weighted 3D gradient echo (3D-GE) sequence acquired on a 7-Tesla scanner. All the data were averaged and resulted in a 40 μm isotropic resolution dataset.

The μ-CT datasets were acquired using the μ-CT module of a MultiModality hybrid micro-Positron Emission Tomography (μ-PET) / μ-CT Inveon Scanner (Siemens Medical Solutions). The Inveon scanner is equipped with a 165-mm x 165-mm X-ray camera and a variable-focus tungsten anode X-ray source operating with a focal spot size of less than 50 μm. The scan consisted of a 30-minute whole-head acquisition over an axial field of view of 44 mm and a transaxial of 88 mm with a resolution of 21.7 μm pixels binned to 43.4 μm. 440 projections were acquired using a 1 mm aluminum filter, a voltage of 80 kV, and a current of 500 μA. The data sets were reconstructed using the Feldkamp algorithm^34^.

The hybrid scanner is equipped with a M2M Biovet module used to monitor continuously vital signs. All rats were monitored continuously throughout the scanning session via a respiration sensor pad and electrocardiogram. The imaging scan consisted of initially placing each rat in an induction chamber using 3-5% isoflurane exposure during 2-3-min until the onset of anesthesia. The animal was then subsequently positioned laterally along the bed palate over a thermistor heating pad in which 2.0% to 3.0% isoflurane was administered via a 90° angled nose cone throughout the scan. The head of each subject was judiciously oriented perpendicular to the axis of the rat body so that the extracranial part of the implanted electrode could be easily kept away from the field of view of the μ-CT image acquisition. Importantly, the large extracranial metal components and dental cement of the implant can cause beam hardening that can appear as cupping, streaks, dark bands or flare in the μ-CT^35–37^. To this effect, the head positioning helped reduce the risks of image artifacts that could be induced by the implant along the path of the X-ray beam.

Locus coeruleus was manually segmented and color-coded with guidance from the Paxinos and Watson Rat Brain Atlas^38^. This region was set as the target of reference. A rigid co-registration between the acquired μ-CT and the modified μ-MRI atlas images was systematically performed using a commercial software Amira (Thermo Fisher Scientific). Both datasets were overlaid to match the intracranial space between both imaging modalities with location of bregma and lambda used as cranial anchors towards stereotaxic localization. Visual analysis helped determine the sub-millimetric localization of the electrode tip. This analysis was conducted by two individuals blind to the behavioral and viral status of the animal. Three animals were excluded from the localization analysis of **Extended Data Fig. 8** as they lacked immunohistochemical data on optical fiber placement; one of those animals had no viable μ-CT signal, and the other two animals were not imaged for μ-CT (including for intracochlear electrode placement).

#### Immunohistochemistry

At the end of the behavioral, imaging, and electrophysiological studies, animals were perfused with 4% paraformaldehyde, brains recovered, and embedded in Optimal Cutting Temperature compound prior to freezing at –80°C. Afterwards, 20 µm thick slices were cut from the brainstem and stained using standard immunohistochemistry histological methods. Staining for tyrosine hydroxylase (primary antibody 1:500, chicken anti-TH, Aves Labs TYH; secondary antibody, 1:1000, goat anti-chicken, Alexa Fluor® 568, Abcam ab175477) was co-localized with YFP (primary antibody 1:500, rabbit anti-GFP, Abcam ab290; secondary antibody 1:1000, goat anti-rabbit, Alexa Fluor® 488, Abcam AB150077).

#### Electrophysiology

ECAP thresholds from the auditory nerve were measured using autoNRT™ (Custom Sound® Suite 4.0, Cochlear Ltd.) an automated system for the Nucleus® Freedom™ cochlear implant (Botros et al., 2007). Traces were analyzed for ECAP magnitude using a custom Matlab script.

For cochlear implant-evoked multi-unit cortical responses, animals were anesthetized as described above. Following partial resection of the temporalis muscle, the temporal skull was removed to expose auditory cortex contralateral to the cochlear implanted ear. In vivo multi-unit recordings were made with a Multiclamp 700B amplifier (Molecular Devices). Recordings were obtained from 500-700 µm below the pial surface with tungsten multi-unit electrodes (0.5 MΩ).

The CI24RE implant emulator was driven by a Freedom system sound processor connected through the Freedom Programming Pod to a Windows personal computer running the Custom Sound EP software (Cochlear Ltd.), and the evoked auditory brainstem response (EABR) function was used (5 charge balanced biphasic pulses, 25 µs/phase, 900 Hz stimulation frequency, 57 μs pulse duration, 20 sweeps at 0.9 Hz) to stimulate the individual electrodes of the array. A modified cable with stereo jack and Bayonet Neill-Concelman connectors was used to connect the Programming Pod to the trigger input of the Digidata 1440A (Molecular Devices), facilitating coordination of each sweep of the electrical stimulus (stimulation intensity was controlled through Custom Sound) with the multi-unit activity recording setup in Clampex 10.3 (Molecular Devices). For cortical analysis, z-scored cochlear implant evoked responses and neural d’ were calculated using a custom Matlab script, quantifying number of spikes in the 100 ms post-stimulus period compared to pre-stimulus baseline. Spike thresholding was based on the RMS of the baseline period. Auditory cortical recording sites with no z-scored evoked activity ≥1 or with spontaneous firing rates ≤ 1 Hz were considered to be non-responsive and not included in analyses. Spike thresholding and spontaneous firing criteria were based on previously published rat auditory cortical firing rates with comparable anesthesia^39^. The neural d’ was computed as the difference in the z-score of the behavioral target channel and the average of the z-scores of the behavioral foil channels. The coefficient of variance was calculated as the standard deviation of z-scored activity for all foil channels divided by the average of the z-scored activity for all foil channels.

#### Statistics

Behavioral data from sham-paired cochlear implant rats in **Figure 1e-h** also served as the control for **Figure 3c-g**. Impedance measurements from sham-paired cochlear implant rats in **Extended Data Figure 5c** was also shown for comparison in **Extended Data Figure 10b**. Paired two-tailed Student’s t-tests were performed in **Figure 1d** and **Extended Data Figure 4a-c**, and unpaired two-tailed Student’s t-tests were performed in **Figure 2e,g**, **Figure 3e**, **Extended Data Figure 7d,e,g,h**, **Extended Data Figure 9b**, and **Extended Data Figure 10c-e**. Unpaired two-tailed Mann-Whitney tests were used in **Figure 3d**, **Extended Data Figure 5b**, **Extended Data Figure 9a**, and **Extended Data Figure 10a**. Pearson’s correlations were computed in **Figure 1i**, **Figure 4g-j**, and **Extended Data Figure 5d-i**. One-way ANOVA with Bonferroni correction was conducted in **Figure 4c**. Error bars and shading on line plots denote ± s.e.m. unless otherwise stated.

### Data availability

The data that support the findings of this study are available from the corresponding author upon request. Source data are provided with this paper.

### Code availability

Source code is available from the corresponding author upon request.

## Acknowledgements

We thank M. Azadpour, N. Capach, I. Carcea, M. Chesler, M. Donegan, P. Gibson, Z. Gironda, M. Insanally, J. Kirk, D. Lin, K.A. Martin, O. Mishkit, J. Multani, J. Neukam, J.T. Roland Jr., E. Sagi, D. Sanes, S. Sara, J.K. Scarpa, J. Schiavo, M. Semerkant, I. Shehu, D. Smyth, J. Tranos, C. Treaba, N. Tritsch, S. Valtcheva, and S. Waltzman for comments, discussions, and technical assistance. We thank Cochlear Ltd. for technical support. We thank the Genotyping Core Laboratory of NYU Langone Health for help with genotyping transgenic rats. We thank the Stanford Neuroscience Gene Vector and Virus Core and the Deisseroth lab for AAVDJ-ef1α-DIO-GCaMP6s (**Figure 2 and Extended Data Figures 6, 7**). We thank C. Schaulsohn for artwork in **Figures 1a, 2a,** and **3b**. This work was funded by a Vilcek Scholar Award (to E.G.); a Hirschl/Weill-Caulier Career Award (to R.C.F.); a Sloan Research Fellowship (to R.C.F.); and the National Institutes of Health (grant number F30-DC017351 to E.G., T32GM007308 to E.G., R01-DC003937 to M.A.S., and R01-DC012557 to R.C.F.). Partial support was also received from a research contract from Cochlear Ltd. to J. Thomas Roland, Jr., M.D. In vivo imaging was performed under the DART Preclinical Imaging Core partially funded by the NYU Laura and Isaac Perlmutter Cancer Center Support Grant, NIH/NCI P30CA016087. The Center for Advanced Imaging Innovation and Research (CAI2R, www.cai2r.net) at NYU School of Medicine is supported by NIH/NIBIB P41 EB017183.

## Author contributions

E.G. conducted electrophysiology, in vivo optogenetics, fiber photometry, and cochlear implant training. E.G. and A.Z. conducted behavioral testing, IHC, and CT/MRI co-registration analysis. All other analysis was done by E.G. Y.Z.W. designed co-registration analysis. E.G, M.A.S., and R.C.F designed the study and wrote the paper.

## Extended Data Figures and Legends

**Extended Data Figure 1.**
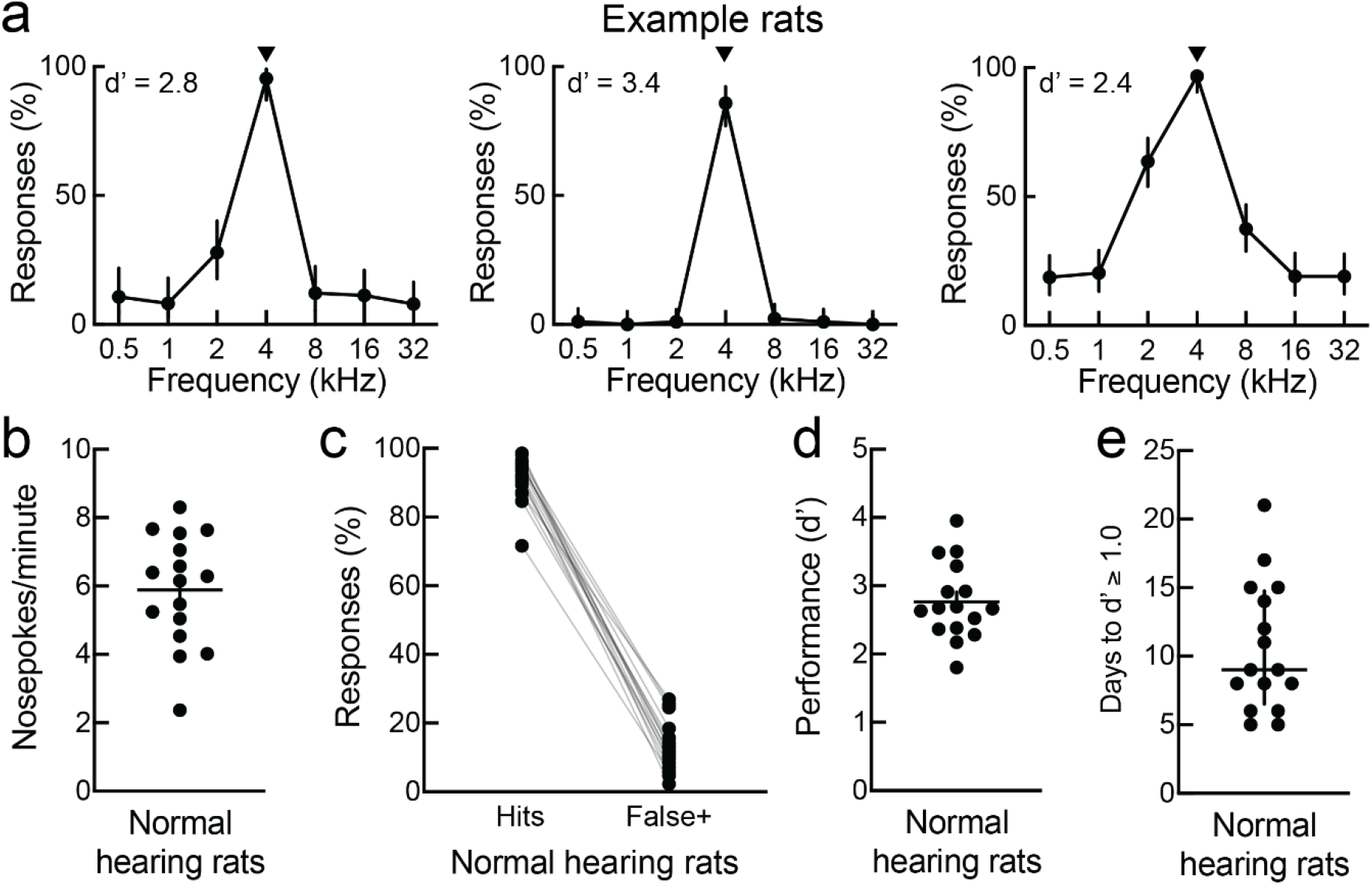
Auditory conditioning on go/no-go task in normal-hearing rats. **a,** Normal-hearing behavioral response curves from three example rats that reached training criteria. Arrowhead, target tone was 4 kHz for all animals. Error bars, 95% confidence intervals. **b,** Average initiation rates for final five days of normal-hearing behavioral performance (N = 16). Error bars, mean ± s.e.m. **c,** Average hits and false positive rates for final five days. **d,** Average behavioral performance (d’) for final five days. Error bars, mean ± s.e.m. **e,** Days to d’ ≥ 1.0. Error bars, median ± interquartile range.

**Extended Data Figure 2.**
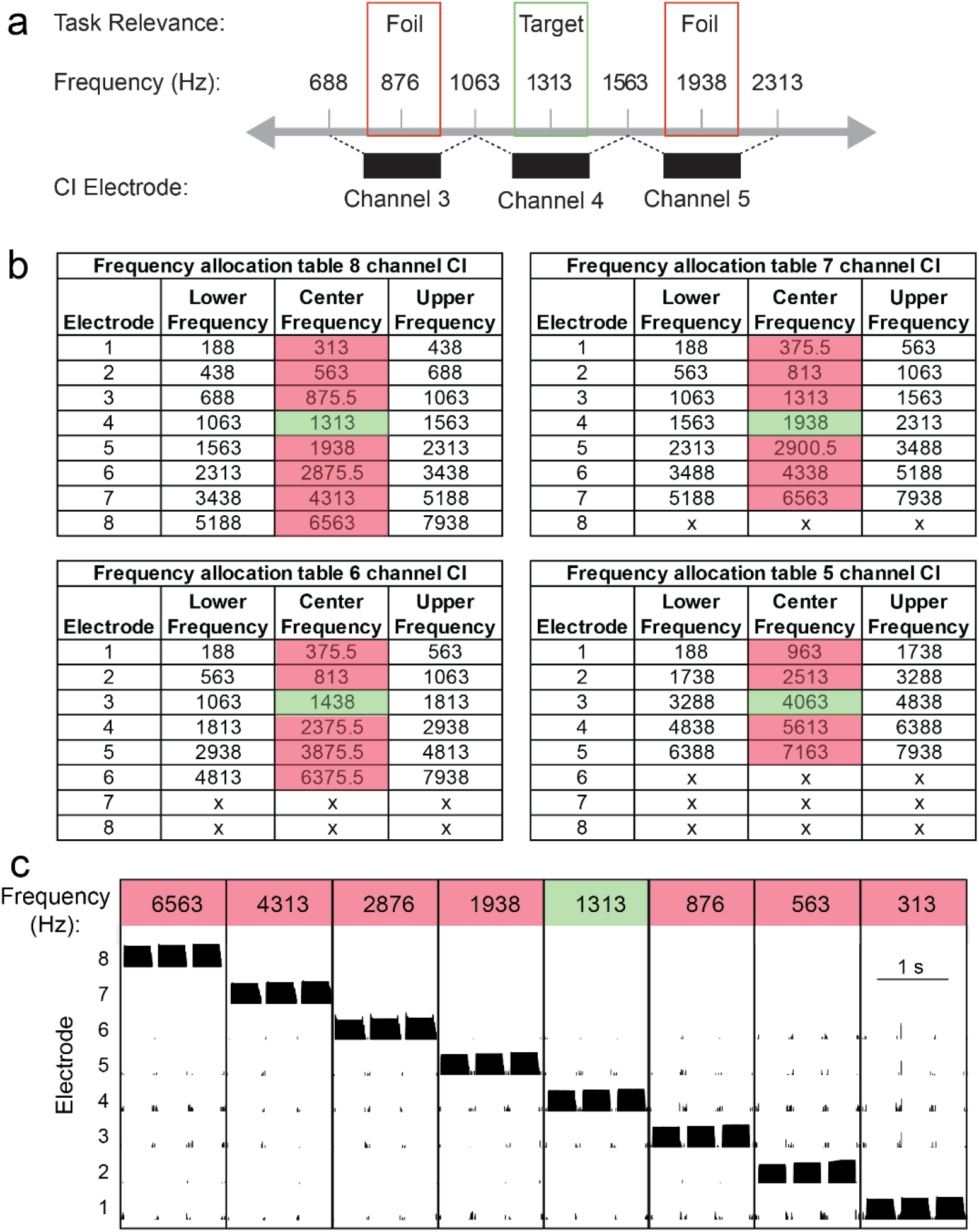
Cochlear implant programming. **a,** Depiction of center frequencies for individual cochlear implant channels. **b,** Frequency allocation tables used to select tones for behavioral conditioning based on the center frequency of channels with different electrode configurations. **c,** Example electrodograms, showing that only the cochlear implant channel for selected center frequency was activated by the tone.

**Extended Data Figure 3.**
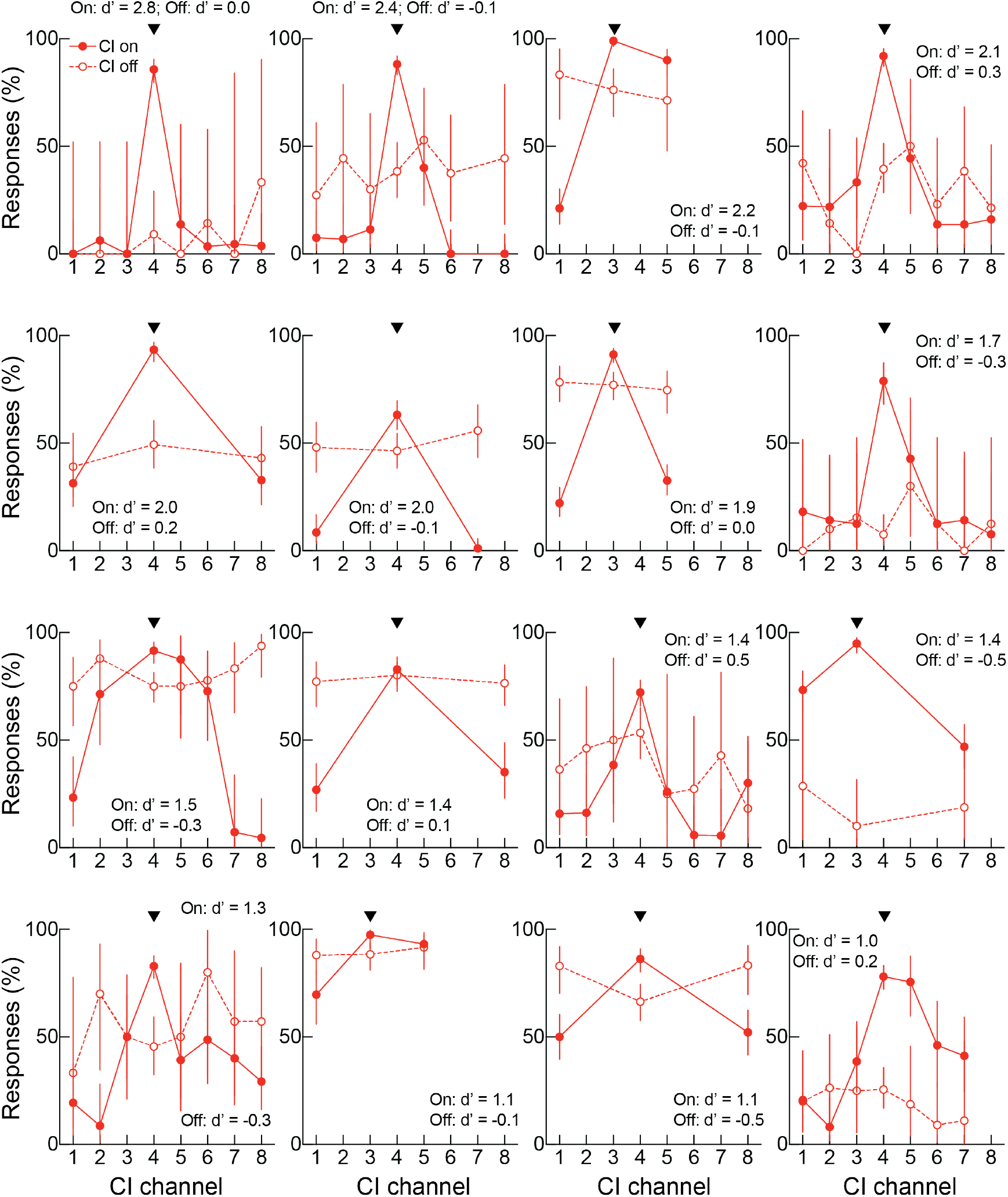
Deafened animals used the cochlear implant to perform the auditory task. Response rates across cochlear implant channels for all 16 rats from Figure 1 with the cochlear implant turned on (filled circles) or turned off (open circles). Arrowhead, target tone programmed to activate channel 3 or 4. Error bars, 95% confidence interval.

**Extended Data Figure 4.**
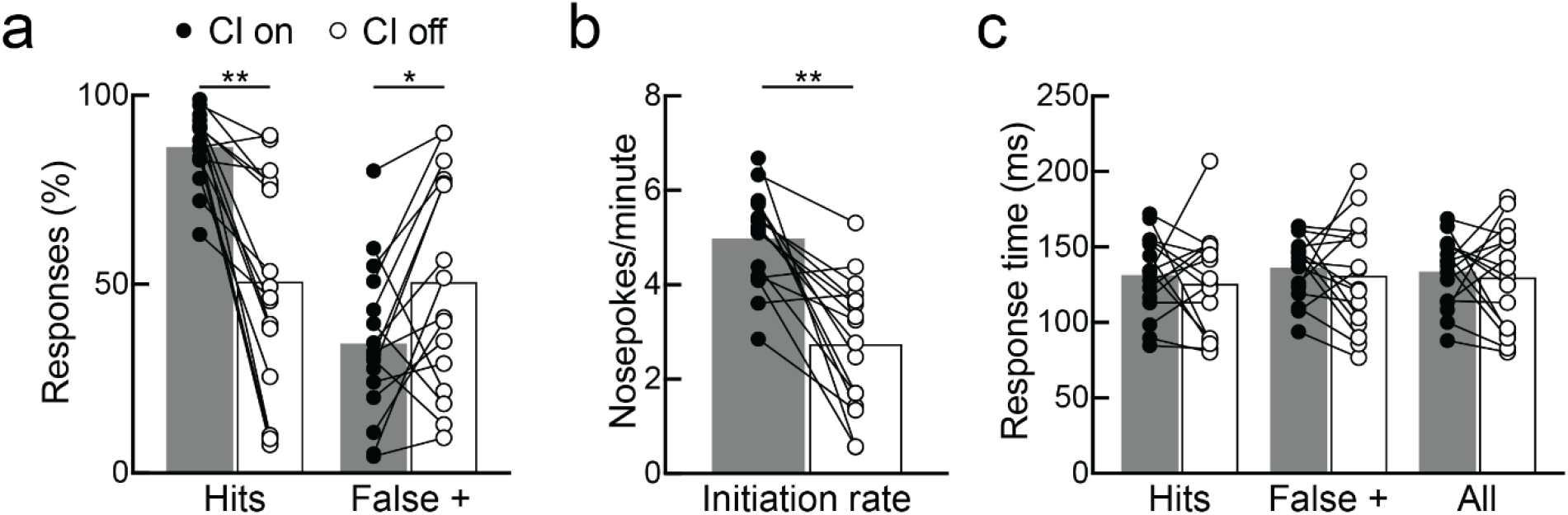
Deafened animals were unable to perform the auditory task without cochlear implant input. **a,** Hit rates were lower and false positives were higher in rats when the cochlear implant was off (N = 16, on vs off, hits: p = 0.0001; false positives: p = 0.02; paired two-tailed Student’s t-tests). **b,** Initiation rates decreased when the cochlear implant was turned off (on vs off, p < 0.0001; paired two-tailed Student’s t-test). **c,** Response times were similar across trial types, whether the cochlear implant was on or off (on vs off, hits: p = 0.58; false positives: p = 0.59; all: p = 0.70; paired two-tailed Student’s t-tests). *, p < 0.05; **, p < 0.01.

**Extended Data Figure 5.**
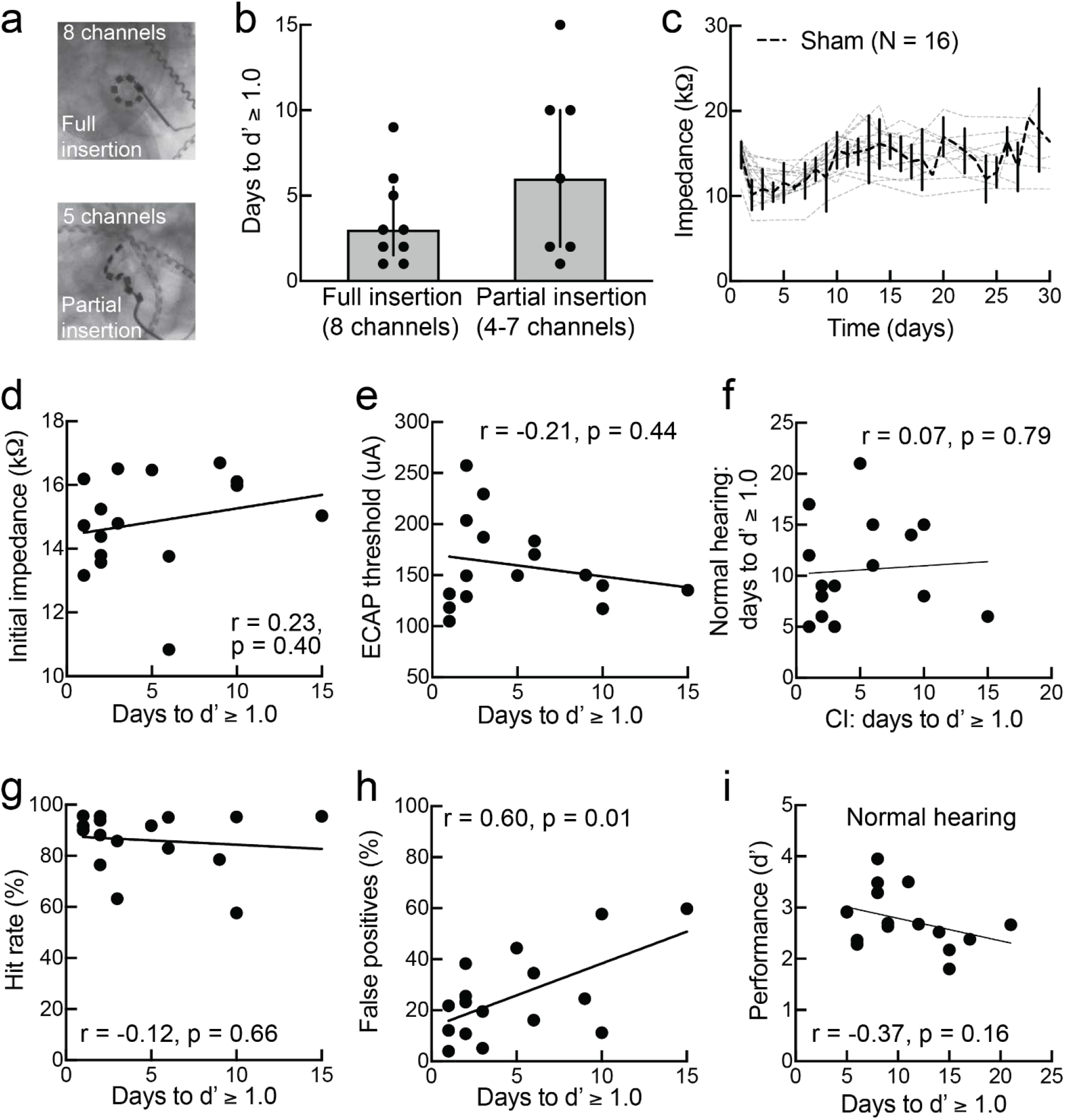
Individual variability with implant use was related to false positive rate but not insertion depth, impedance, ECAP thresholds, hit rates, or normal-hearing performance. **a,** Example x-rays of full insertion (8 channels) and partial insertion (4-7 channels). **b,** Days to d’ ≥ 1.0 did not differ based on cochlear implant insertion depth (full insertion vs partial insertion, p = 0.32, unpaired two-tailed Mann–Whitney test). Median ± interquartile range. **c,** Average impedance of active cochlear implant channels over time. Grey dashed lines, individual rats. Black, mean ± s.e.m. **d,** Days to d’ ≥ 1.0 did not correlate with initial impedance values (N = 16, Pearson’s r = 0.23, p = 0.40). **e,** Days to d’ ≥ 1.0 did not correlate with ECAP threshold (Pearson’s r = −0.21, p = 0.44). **f,** Cochlear implant learning days to d’ ≥ 1.0 did not correlate normal-hearing learning days to d’ ≥ 1.0 (Pearson’s r = 0.07, p = 0.79). **g,** Days to d’ ≥ 1.0 did not correlate with hit rate (Pearson’s r = 0.16, p = 0.56). **h,** Days to d’ ≥ 1.0 correlated with false positives (Pearson’s r = 0.61, p = 0.01). **i,** During normal-hearing training, days to d’ ≥ 1.0 did not correlate with maximum d’ performance (Pearson’s r = −0.37, p = 0.16).

**Extended Data Figure 6.**
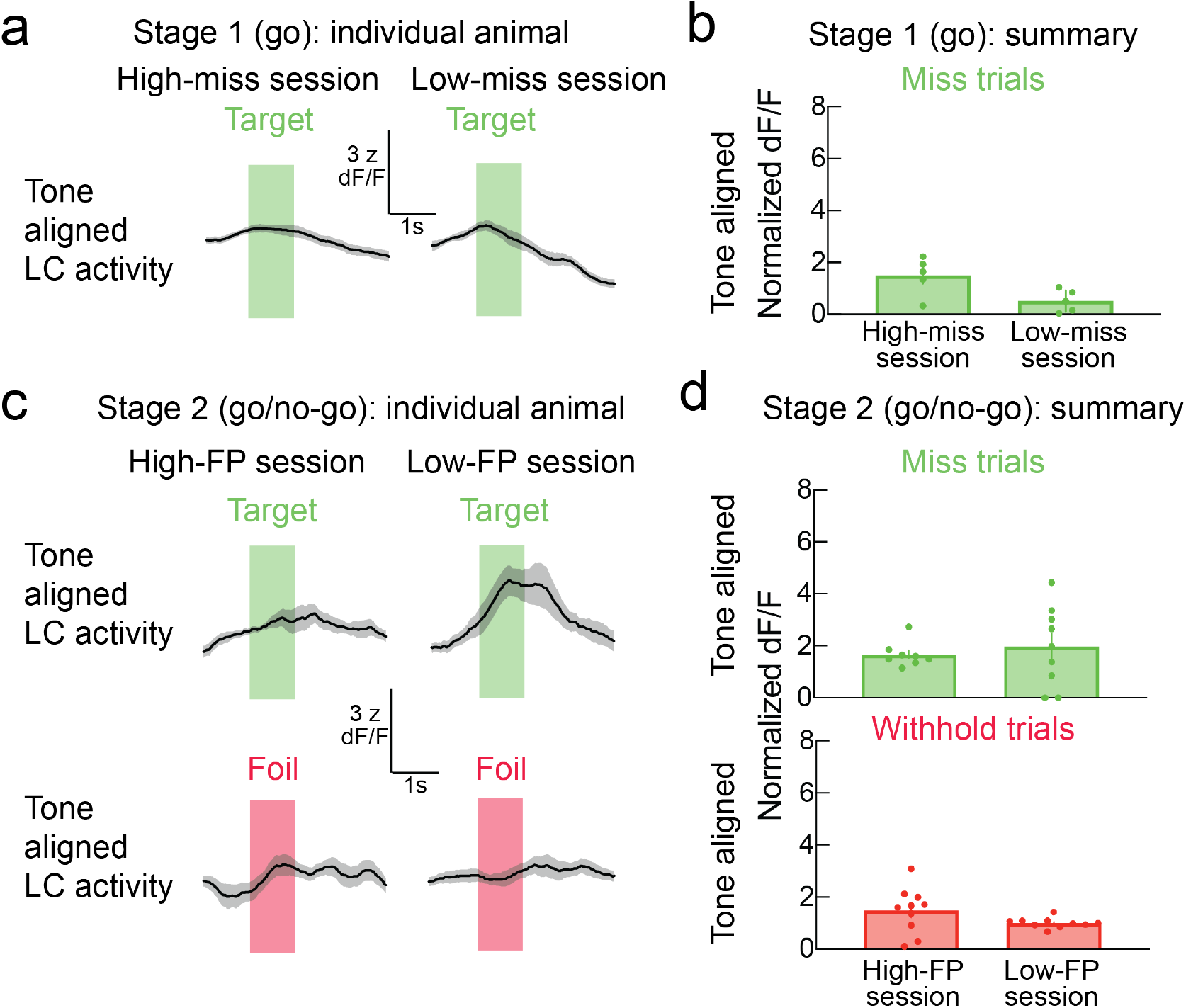
Fiber photometry miss/withhold analysis. **a,** Example locus coeruleus (LC) activity aligned to tone onset during stage one training with cochlear implant, showing trial-averaged misses in an early high-miss rate behavioral session (left) and in a later low-miss behavioral session (right). **b,** Tone-aligned normalized dF/F locus coeruleus signals in high-miss and low-miss behavioral sessions. **c,** Example locus coeruleus activity aligned to tone onset during stage two (foil and target training). Miss and withhold trials in a high-false positive (FP) behavioral session (left). Miss and withhold trials in a low-FP behavioral session (right). **d,** Tone-aligned normalized dF/F locus coeruleus signals during miss trials and withhold trials in high-FP and low-FP behavioral sessions. Data are mean ± s.e.m.

**Extended Data Figure 7.**
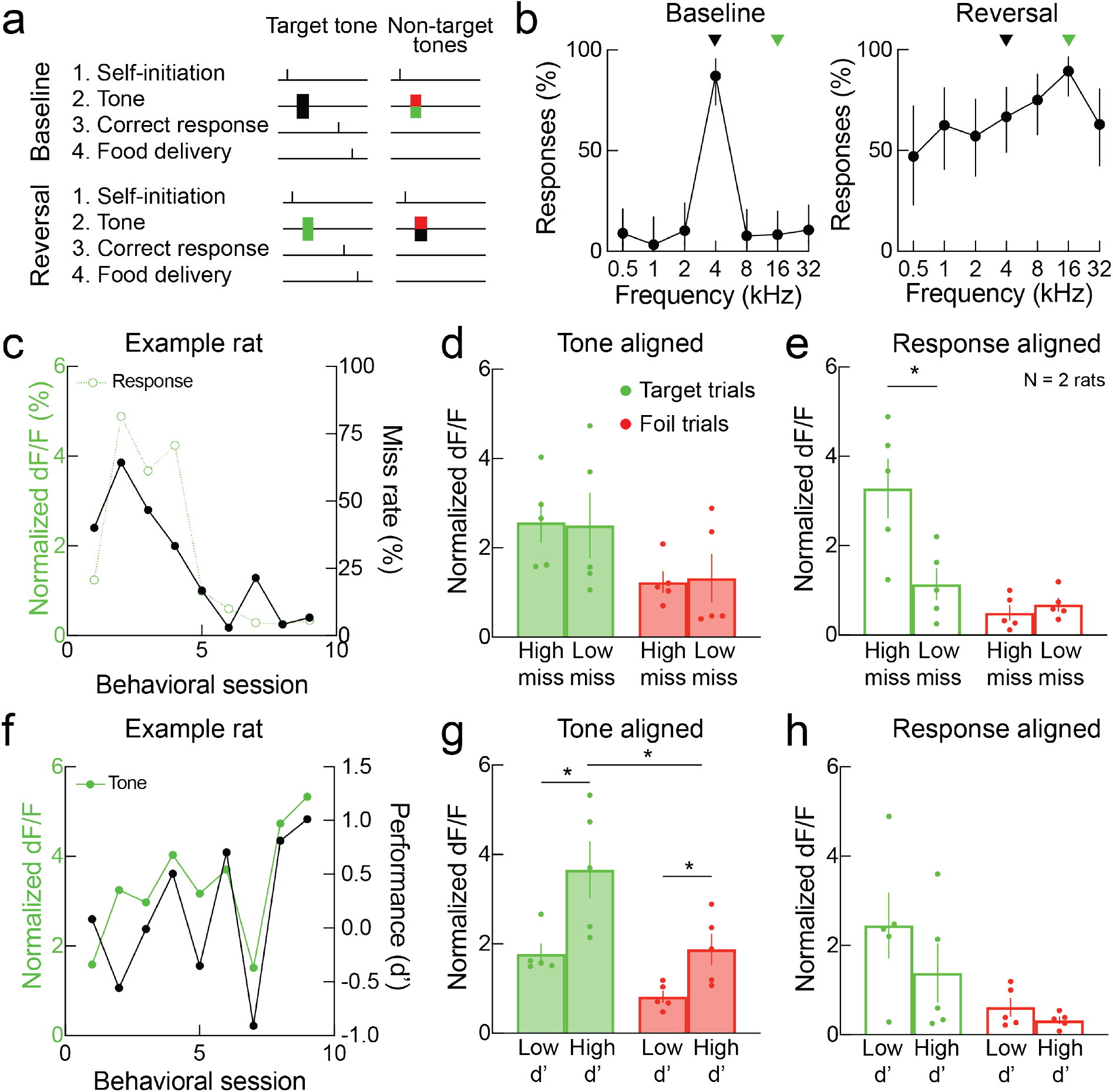
Locus coeruleus activity in normal-hearing reversal learning. **a,** Schematic of go/no-go auditory behavioral task in normal-hearing rats when target tone is changed to a different frequency. After training to response to one target tone (black) while withholding from foil tones (green/red), one of the previously unrewarded tones (green) became the rewarded tone and the previously rewarded tone (black) became unrewarded. **b,** Example of animal performance on this task to first and second rewarded tones. Black arrowhead, first rewarded tone; green arrowhead, second rewarded tone. Error bars, 95% confidence intervals. **c**, Example of response-aligned locus coeruleus activity and miss rates across time. **d,** Tone-aligned normalized dF/F in locus coeruleus was similar across miss rate sessions on hit target trials (p = 0.93, unpaired two-tailed Student’s t-test) and false positive foil trials (p = 0.88, unpaired two-tailed Student’s t-test). **e,** Response-aligned normalized dF/F in locus coeruleus was higher on target hit trials in high-miss sessions (p = 0.02, unpaired two-tailed Student’s t-test), but comparable across miss rates for false positive foil trials (p = 0.43, unpaired two-tailed Student’s t-test). **f,** Example tone-aligned locus coeruleus activity and performance (d’) across time. **g,** Tone-aligned normalized dF/F in locus coeruleus was higher when d’ was high on hit target trials (p = 0.02, two-tailed unpaired Student’s t-test), as well as on false positive foil trials (p = 0.02, unpaired two-tailed Student’s t-test), but higher on hit trials than on foil trials (p = 0.04, unpaired two-tailed Student’s t-test). **h,** Response-aligned normalized dF/F in locus coeruleus was similar across d’ values on target hit trials (p = 0.31, unpaired two-tailed Student’s t-test) and for false positive foil trials (p = 0.21, unpaired two-tailed Student’s t-test). Data are mean ± s.e.m. *, p < 0.05.

**Extended Data Figure 8.**
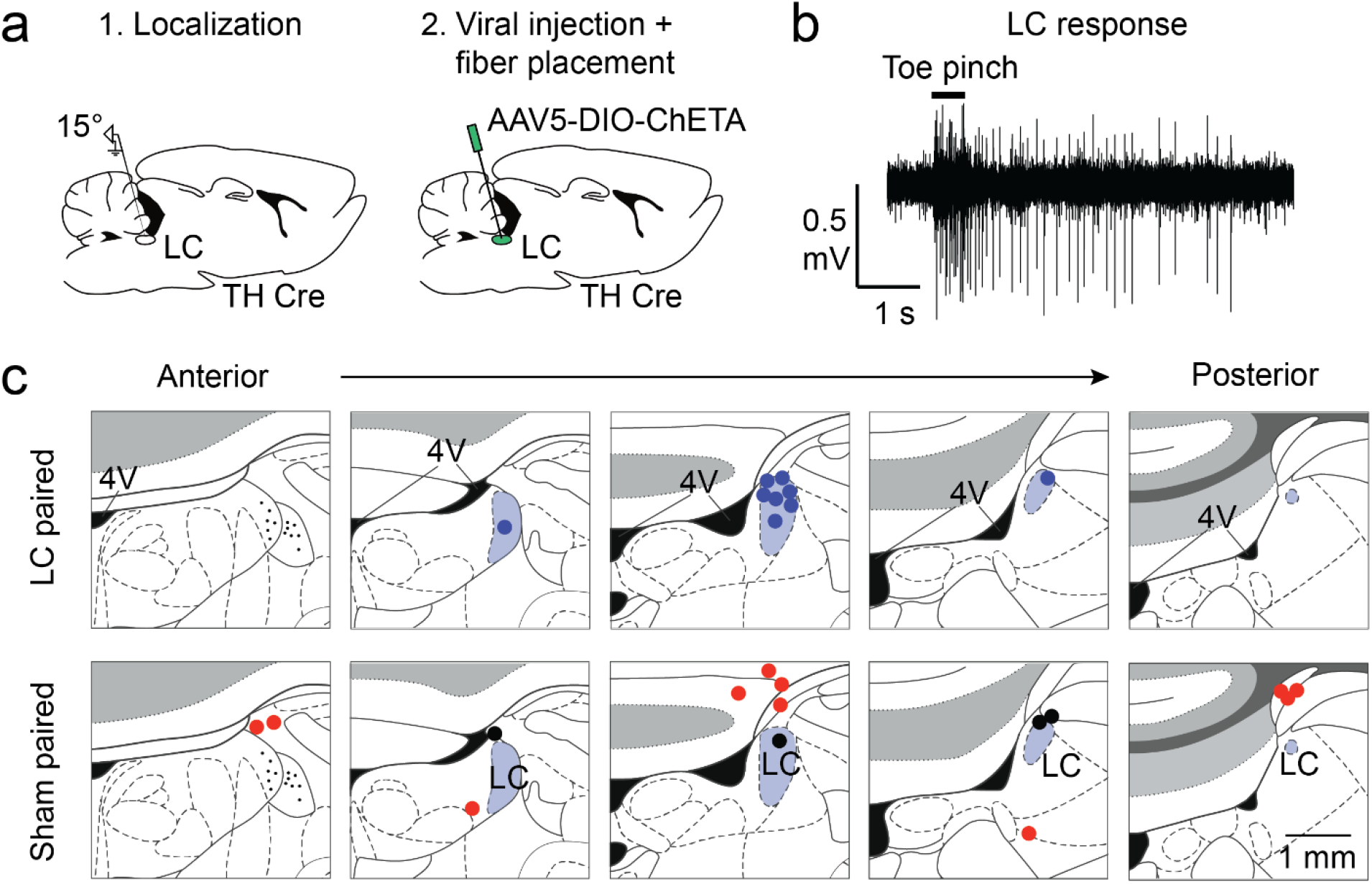
Locus coeruleus targeting. **a,** Surgical approach for targeting locus coeruleus. Multi-unit recordings were conducted to locate locus coeruleus (LC) and then viral injection and optic fiber placement were based on these coordinates. **b,** Example multi-unit activity in locus coeruleus evoked by toe pinch. **c,** Optical fiber placement based on histology and μ-CT/ μ-MRI co-registration. Top, fiber placement in locus coeruleus paired animals. Bottom, fiber placement in sham animals (red, mistargeted fibers outside of locus coeruleus; black, YFP-injected controls). Scale bar, 1 mm.

**Extended Data Figure 9.**
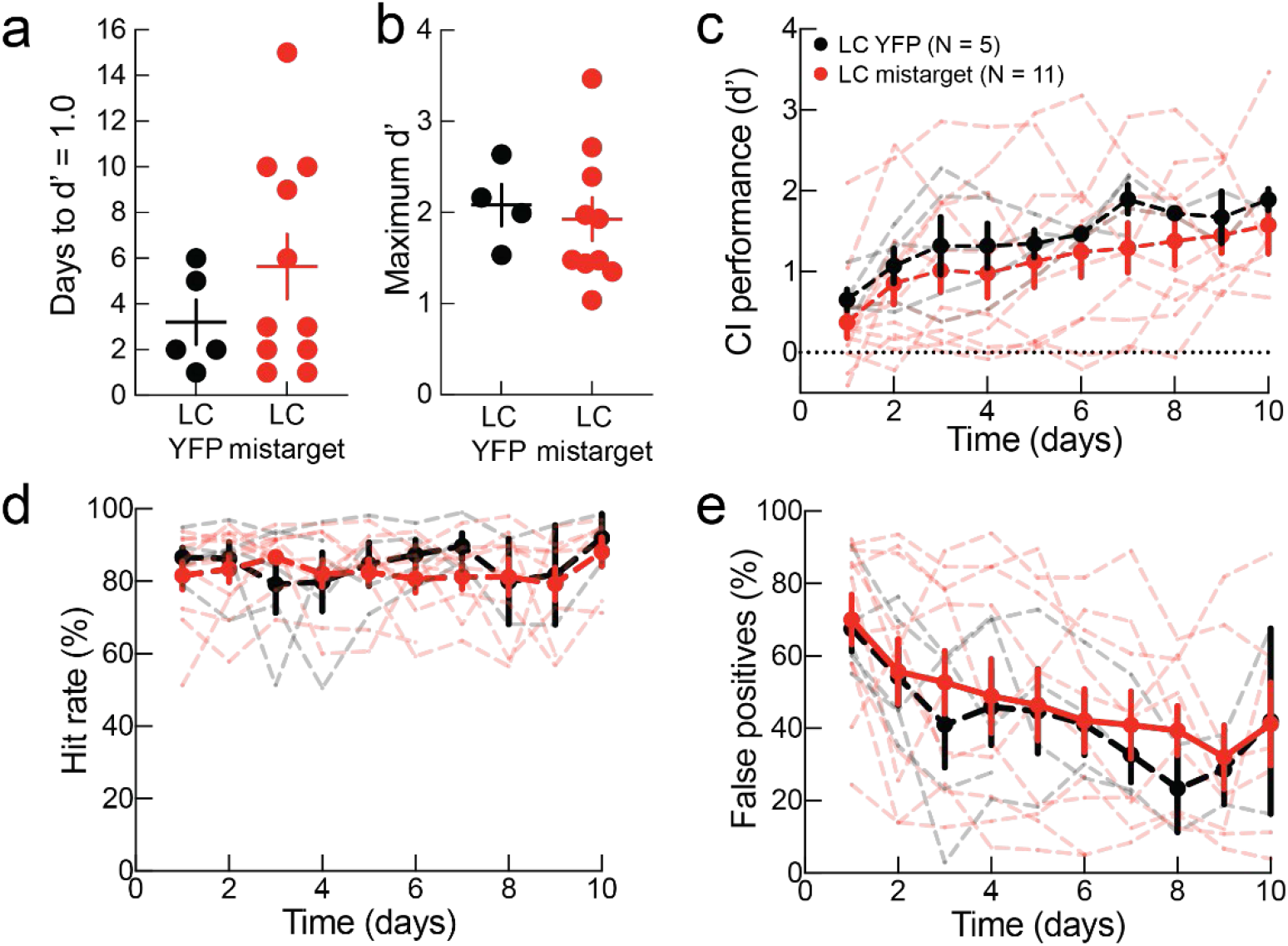
Cochlear implant performance was comparable between sham paired YFP-injected animals vs fiber-mistargeted animals. **a,** Days to d’ ≥ 1.0 was similar between the two sub-groups of sham animals with either YFP-only expression in locus coeruleus (LC) or when fiber was mistargeted outside of locus coeruleus (YFP: N = 5 rats, mistargeted: N = 11 rats, p = 0.38, unpaired two-tailed Mann–Whitney test). Median ± interquartile range. **b,** Sham animals in each subgroup with at least six days of cochlear implant training had similar maximum d’ (YFP: N = 4 rats vs, mistargeted: N = 10 rats, p = 0.71, unpaired two-tailed Student’s t-test). **c,** Implant performance (d’) over time in YFP vs mistargeted animals. One YFP animal and one mistargeted animal shown in **a,c** each did not reach the six-day requirement for maximum performance analysis; these animals are not displayed in **b**. **d,** Hit rates. **e,** False positives. Data are error bars, mean ± s.e.m. except in **a**.

**Extended Data Figure 10.**
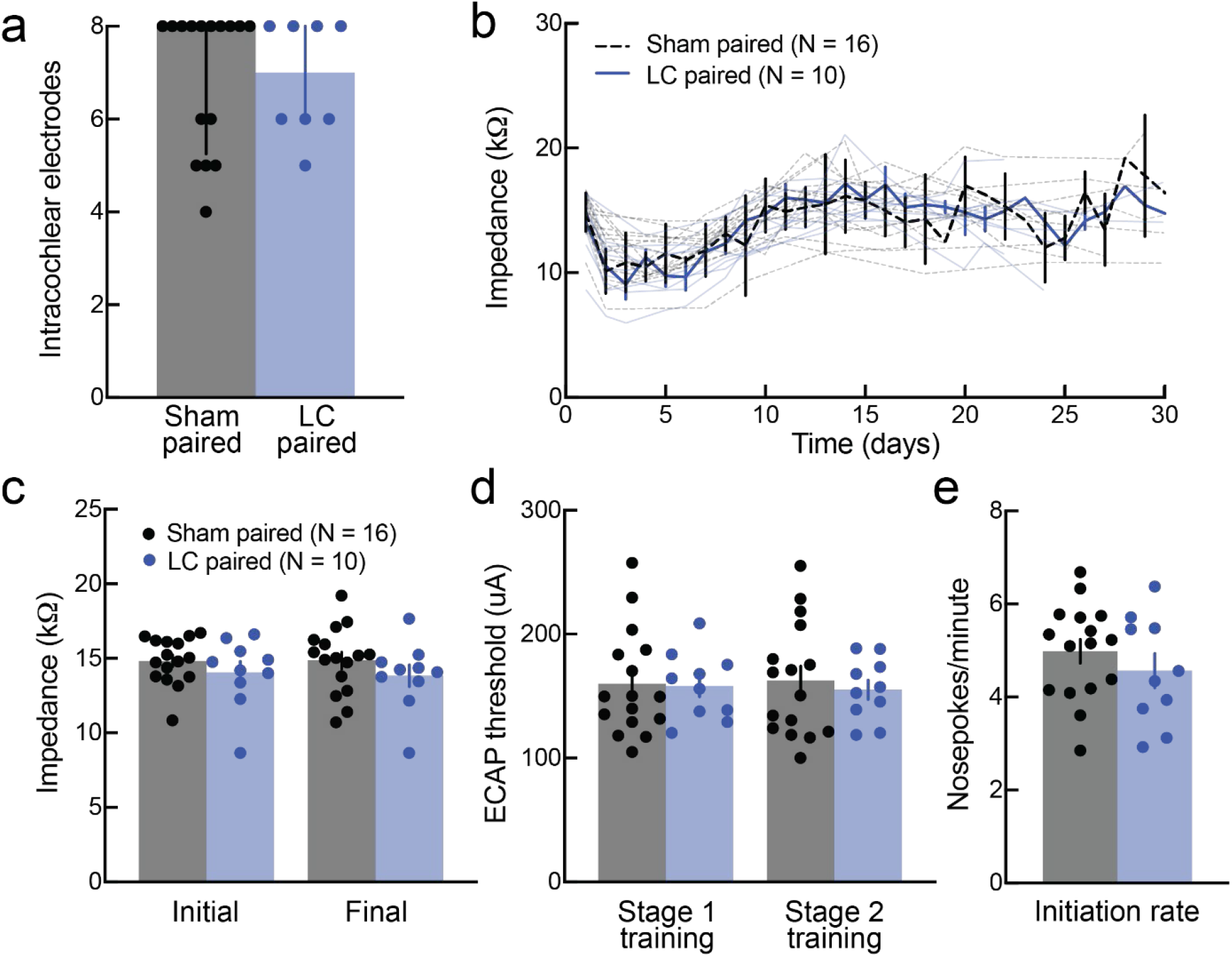
Locus coeruleus vs sham paired animals had comparable implant insertions, impedances, ECAPs, behavioral initiation rates, and lack of residual hearing. **a,** Number of intracochlear electrodes as assessed by x-ray was similar between locus coeruleus (LC) paired rats and sham paired rats (locus coeruleus paired, N = 10 rats vs sham paired, N = 16 rats, p = 0.93, unpaired two-tailed Mann–Whitney test). Error bars, median ± interquartile range. **b,** Average impedances of cochlear implant channels over time in locus coeruleus paired vs sham paired rats. **c,** Both initial and final impedance values were similar in locus coeruleus paired and sham paired rat (locus coeruleus paired vs sham paired, initial: p = 0.32; final: p = 0.27; unpaired two-tailed Student’s t-test). **d,** ECAP thresholds during stage one and stage two training did not differ between locus coeruleus paired and sham paired rats (locus coeruleus paired vs sham paired, stage one: p = 0.91; stage two: p = 0.64; unpaired two-tailed Student’s t-test). **e,** Initiation rates were similar between locus coeruleus paired and sham paired rats (locus coeruleus paired vs sham paired, p = 0.35; unpaired two-tailed Student’s t-test). Data are error bars, mean ± s.e.m. except in **a**.

## Supplementary Movies

**Movie S1**

Example of profoundly deaf rat using cochlear implant to respond to presentation of target tones but not foil tones.

